# Orthogonal Gene Engineering Enables CD8^+^ T Cells to Control Tumors through a Novel PD-1^+^TOX-indifferent Synthetic Effector State

**DOI:** 10.1101/2022.02.18.481059

**Authors:** Jesus Corria-Osorio, Santiago J. Carmona, Evangelos Stefanidis, Massimo Andreatta, Tania Muller, Yaquelin Ortiz-Miranda, Bili Seijo, Wilson Castro, Cristina Jimenez-Luna, Leonardo Scarpellino, Catherine Ronet, Aodrenn Spill, Evripidis Lanitis, Sanjiv A. Luther, Pedro Romero, Melita Irving, George Coukos

## Abstract

Adoptive immunotherapy offers opportunities to reprogram T cells and the tumor microenvironment. Orthogonal engineering of adoptively transferred T cells with an IL-2Rβγ-binding IL-2 variant, PD1-decoy and IL-33 led to cell-autonomous T-cell expansion, T-cell engraftment and tumor control in immunocompetent hosts through reprogramming of both transferred and endogenous CD8^+^ cells. Tumor-infiltrating lymphocytes adopted a novel effector state characterized by TOX suppression and specific expression of multiple effector molecules, most prominently granzyme C. While the IL-2 variant promoted CD8^+^ T-cell stemness and persistence, and was associated with downregulation of TOX, the combination with IL-33 was necessary to trigger the novel polyfunctional effector state. Rational T-cell engineering without host lymphodepletion enables optimal reprogramming of adoptively transferred T cells as well as mobilization of endogenous immunity into new functional CD8^+^ states mediating tumor control.

## Introduction

Adoptive T-cell therapy (ACT) has emerged as a powerful approach against advanced human tumors (June, Riddell, and Schumacher 2015; Klebanoff, Rosenberg, and Restifo 2016). Expansion and persistence of adoptively transferred T cells in recipient patients are required for effective ACT. Host preconditioning with non-myeloablative chemotherapy and post-ACT support with high-dose IL-2 (Rosenberg and Restifo 2015) have been commonly used to enable engraftment, expansion and persistence of adoptively transferred autologous T cells, but are associated with important toxicity and clinical costs (Wolf et al. 2019). Use of T-cell subsets with stem-like potential or T-cell conditioning *in vitro* during culture have been proposed as approaches to enhance cell persistence and the efficacy of ACT (Vodnala et al. 2019; Crompton, Sukumar, and Restifo 2014). The modification of T cells with synthetic costimulatory modules has also enabled remarkable tumor responses upon ACT. However, significant barriers in the tumor microenvironment (TME) still limit T-cell engraftment, persistence and function in solid tumors (Anderson, Stromnes, and Greenberg 2017), and T-cell exhaustion remains an important reason of ACT failure (Lynn et al. 2019; Feucht et al. 2019). The use of T-cell precursors, reprogramming cultures, host lymphodepletion, IL-2 support, costimulatory signaling, or blockade of programmed cell death-1 (PD-1) or its ligand-1 (PD-L1) have been used to ensure that transferred T cells expand sufficiently *in vivo* and acquire a functional state mediating tumor rejection. However, the precise nature of such desired state(s) remains elusive.

We hypothesized that T cells could be rewired to both enable their engraftment in the absence of lymphodepletion and confer a functional state able of rejection of poorly immunogenic tumors. We sought to engineer T cells using an orthogonal combinatorial approach to reprogram them, and enable them to reprogram endogenous immunity in the TME. We targeted the PD-1/PD-L1 inhibitory pathway with a secreted PD-1 decoy (PD1d), i.e. a fusion molecule comprising the ectodomain of murine PD-1 linked to the Fc region of human IgG4. We also introduced a human IL-2 variant (IL-2^V^), which does not engage the high-affinity IL-2 receptor α-chain (CD25), exhibits decreased toxicity and stimulates selectively effector lymphocytes over regulatory T cells (Carmenate et al. 2013; Rojas et al. 2019). Since the interaction of IL-2 with CD25 favors the formation of more terminally differentiated T cell subsets (Kalia et al. 2010; Obar et al. 2010; Pipkin et al. 2010), we hypothesized that IL-2^V^could promote CD8^+^T-cell stemness, a favorable feature for ACT (Chan et al. 2021). Finally, we introduced IL-33, an alarmin demonstrated to inflame tumors (Villarreal et al. 2014; Kallert et al. 2017), unleash the cross-priming potential of tumor-associated DCs (Gao et al. 2013; Dominguez et al. 2017; Gao et al. 2015; Villarreal and Weiner 2014), and favor CD8^+^ T-cell immunity (Villarreal et al. 2014; Kallert et al. 2017). As proof of principle, we orthogonally engineered OT1 lymphocytes to treat advanced, poorly immunogenic B16-OVA melanoma tumors (Yu et al. 2018) in immunocompetent recipient mice. Here, we characterize in great detail the resulting CD8^+^ T-cell states required for achieving T-cell engraftment and tumor regression in the absence of preconditioning lymphodepletion or exogenous cytokine support.

## Results

### Orthogonal T-cell engineering enables ACT efficacy without lymphodepletion

To begin, we evaluated the PD-1 decoy engineering module. PD1d was secreted efficiently by engineered OT1 cells, and demonstrated specificity of binding to PD-L1 *in vitro* (**Figure S1A**). ACT using PD1d-engineered OT1 cells in lymphodepleted mice bearing small (∼30 mm) tumors, delayed tumor growth (**Figure S1B**). Furthermore, we ascertained that OT1 cells transduced with the PD1d/2^V^ or PD1d/33 modules could secrete simultaneously the two transgenes (PD1d: 50-100 ng/mL plus IL-2^V^: 100-300 ng/mL or IL-33: 10-20 ng/mL per 10^6^ cells over 72 hours, respectively) (**Figure S1C-D**).

We next conducted ACT in mice with more advanced (100 m^3^) tumors in the absence of preconditioning lymphodepletion or exogenous cytokines (**Figure 1A**). Infused OT1 cells in all conditions had mostly a central memory-like phenotype (**Figure S1E**). Two infusions of 5×10^6^ untransduced OT1 cells had minimal impact on tumor growth (**Figure 1B**, **Figure S1F-G**). OT1 cells transduced with PD1d had a minimal effect, and systemic administration of anti-(α)PD-L1 antibody with untransduced OT1 cells produced comparable results to PD1d-OT1 cells. IL-2^V^-engineered or double PD1d/2^V^-engineered OT1 cells did not prove more effective. OT1 cells transduced with IL-33 had minimal but significant effect, as did PD1d/33-OT1 cells. Strikingly, PD1d/2^V^/33-OT1 cells (i.e. 1:1 mixed PD1d/2^V^ and PD1d/33 cells) induced marked tumor regression and achieved an objective tumor response rate (ORR) of 85.7% (predicted probability of occurrence of 83.3%; **Table S1 and S2)**, while ORR was between 0 and 9% for any other treatment. Furthermore, earlier treatment (starting on day 6) led to B16-OVA tumor eradication and cures **(Figure S1H).** In addition, triple-engineered, orthogonal ACT significantly delayed tumor growth and increased survival of mice bearing advanced MC38-OVA colon tumors (**Figure S1I**).

**Figure 1.**
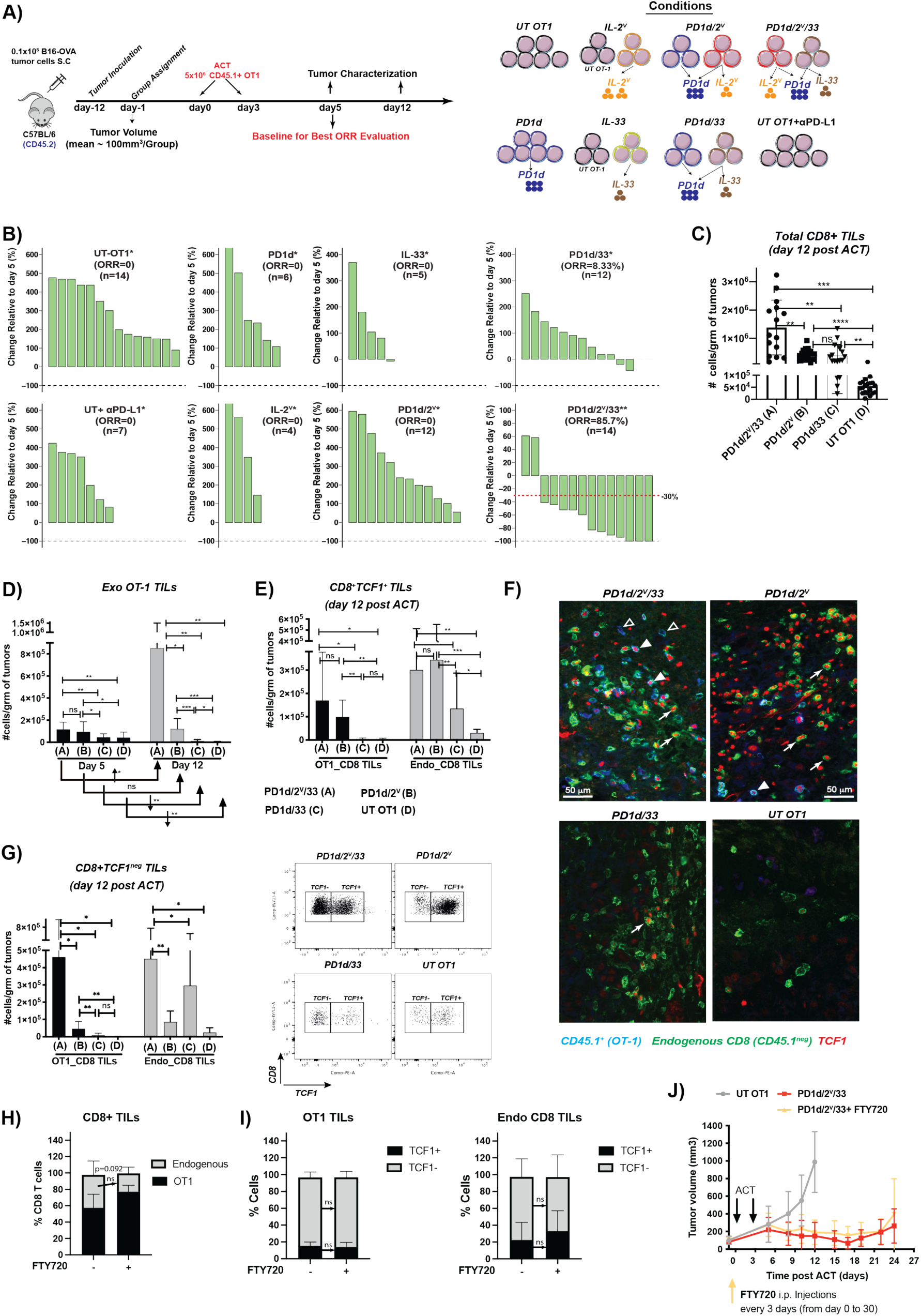
Orthogonal T-cell engineering enables ACT efficacy without lymphodepletion through *in situ* expansion of TCF1^+^ precursor and TCF1^neg^ effector CD8^+^ T cells. **(A)** Experimental design. **(B)** Waterfall plot showing changes in tumor volumes relative to day 5 post ACT. The best response (smallest tumor volume) observed for each animal after at least 12 days post-1^st^ ACT was taken for the calculation (* day 12 post tumor inoculation, ** day 19 post tumor inoculation). Objective Response rate (ORR) includes Complete Response (CR; 100% reduction in tumor volume) and Partial Response (PR; ≤-30% tumor change). **(C-F)** Mice with B16.OVA tumors were treated with either engineered or untransduced OT1 cells as indicated; then tumors were harvested on days 5 and 12 post ACT and cell quantification was performed by flow cytometry. Data are from three independent experiments (n >= 5 animals/group). **(C)** Total numbers of CD8^+^ TILs at day 12. **(D)** Total number of CD45.1^+^ OT1 on days 5 and 12. **(E)** Total numbers of endogenous and exogenous TCF1^+^ CD8^+^ TILs on day 12. **(F)** Representative immunofluorescence micrographs of tumor sections from each experimental group on day 12 showing OT1 and endogenous TCF1^+^ CD8^+^ TILs. Filled triangle: TCF1^+^OT1, open triangle: TCF1^neg^OT1, white arrows: TCF1^+^ Endogenous CD8^+^ TILs. **(G)** Left: total number of OT1 and endogenous TCF1^neg^ CD8^+^ TILs. Data are from three independent experiments (n >= 5 animals/group). Right: a representative dot plot showing the distribution of CD8^+^ TILs of each indicated treatment based on the expression of TCF1. **(H)** Composition of the intratumoral CD8 TIL compartment (Endogenous vs OT1 TILs) 12 days post orthogonal ACT in the presence or absence of FTY720. **(I)** Fraction of TCF1^+^ and TCF1^neg^ in both Endogenous and OT1 TILs harvested 12 days post orthogonal ACT in the presence or absence of FTY720. **(J)** PD1d/2^V^/33+ OT-1 cells were administrated as previously indicated, to B16.OVA-tumor bearing wtC57BL6 mice that also were treated with 100*μ*g/mouse of the drug FTY720 (administrated i.p. every three days beginning two days before 1^st^ cell transfer). A Brown-Forsythe and Welch ANOVA test combined with Tukey Test to correct for multiple comparisons was used for comparing different groups in (C-G) and tumor volumes in (**J**). A two-tailed Student’s t test with Welch’s correction was used for comparing day 5 and day 12 in (D and F) and in (H-I). * p<0.05, ** p<0.01, *** p<0.001, ****p<0.0001.

Orthogonal ACT led to markedly increased T-cell accumulation in tumors. Already by day 5 post ACT, we observed >800-fold more CD8^+^ TILs in tumors treated with PD1d/2^V^/33-ACT relative to baseline (non-treated) B16-OVA tumors (**Figure S2A**). Notably, 12 days post ACT (upon tumor regression) tumors treated with PD1d/2^V^/33-ACT exhibited significantly more total CD8^+^ TILs relative to any other condition (**Figure 1C**). Furthermore, transferred OT1 (CD45.1^+^) cells underwent marked intratumoral expansion specifically following PD1d/2^V^/33-ACT, while PD1d/2^V^-OT1 or PD1d/33-OT1 cells showed modest or minimal levels of tumor expansion, respectively (**Figure 1D, Figure S2B**). Thus, triple-engineering ACT was uniquely associated with marked cell-autonomous expansion of adoptively transferred T cells in tumors, and tumor regression.

### Orthogonal ACT expands TCF1^+^ precursor and TCF1^-^ effector CD8^+^ T cells *in situ*

In conventional ACT schemes, lymphodepletion is required to make available the host homeostatic cytokines thus promoting survival and expansion of the transferred cells (Gattinoni et al. 2005; Muranski et al. 2006). We asked how orthogonally-engineered transferred cells achieved such expansion in the absence of lymphodepletion. TCF1 is a transcription factor essential for early T-cell self-renewal, and TCF1^+^ precursor T cells are required for response to checkpoint blockade immunotherapy (Zhao, Shan, and Xue 2021) and the efficacy of ACT (Chan et al. 2021). We noticed a marked expansion of TCF1^+^ OT1 cells (as well as TCF1^+^ endogenous TILs) when IL-2^V^ was included in the ACT, i.e. with PD1d/2^V^/33- or PD1d/2^V^-engineered OT1 cells (**Figures 1E-F**, **S2C**), suggesting that IL-2^V^ promoted stemness and expansion of transferred cells, thus obviating the need for endogenous homeostatic cytokines associated with lymphodepletion. This effect extended also to the endogenous CD8^+^ TILs. Approximately 10% of OT1 and 30-50% of endogenous CD8^+^ TILs expressed TCF1 post PD1d/2^V^/33-ACT, and the number of total OT1 as well as endogenous CD8^+^ TILs strongly correlated with the number of TCF1^+^ cells in each compartment (**Figure S2D-E**).

Differentiation to effector T cells is associated with downregulation of TCF1 (Danilo et al. 2018; Chen, Ji, et al. 2019). Unsurprisingly, effective tumor control by gene-engineered OT1 cells required high numbers of TCF1^neg^ effector CD8^+^ TILs, a condition met solely following PD1d/2^V^/33-ACT (**Figure 1G**). Indeed, in these tumors 80-90% of OT1 TILs and 50-70% of endogenous CD8^+^ TILs were TCF1^neg^ (**Figure S2C**). Conversely, there were far fewer TCF1^neg^CD8^+^ TILs post PD1d/2^V^-ACT (**Figure 1G**), suggesting that although IL-2^V^ could expand TCF1^+^ precursors, these cells were unlikely to transition to a TCF1^neg^ effector-like state in the absence of IL-33 coexpression. In fact, PD1d/33-ACT was associated with high frequency of TCF1^neg^CD8^+^ TIL, but overall poor TIL expansion and, consistent with the absence of IL-2^V^ from the mix, low TCF1^+^CD8^+^ numbers (**Figure 1E-G**). Remarkably, the important expansion of TCF1^+^ and TCF1^neg^ CD8^+^ TIL observed with orthogonal ACT did not depend on tumor-draining lymph nodes (TDLNs). Indeed, impairment of lymphocyte egress from the lymph nodes by co-administration of FTY720 did not compromise the intratumoral accumulation of OT1 or endogenous CD8^+^ TILs (**Figure 1H, S2F**), or the fraction of TCF1^+^ cells in each compartment (**Figure 1I, S2G**), and did not impact significantly the antitumor efficacy of orthogonal ACT (**Figure 1J**), suggesting that expansion of TCF1^+^ precursors, and subsequent generation of TCF1^neg^ effector CD8^+^ TILs, occurred largely *in situ* thanks to the cooperation of the two cytokines in the TME.

### Orthogonal engineering induces a novel effector CD8^+^ T-cell state

We asked what properties conferred orthogonally engineered cells the ability to achieve regression of non-immunogenic B16 tumors. To this end, we analyzed CD8*^+^* TILs from different ACT settings using single-cell RNA-sequencing (scRNAseq) (**Figure 2A**). Unsupervised clustering analysis of bulk CD8*^+^* TILs revealed five distinct states (clusters C1-C5, **Figure 2B**), which largely reflected the different engineering strategies. TILs from ACT using untransduced T cells clustered in C1, characterized by naïve/central-memory genes (e.g. *Sell*, *Il7r*, *Tcf7*, *Lef1*); and C4, characterized by canonical T-cell exhaustion markers (*Tox*, *Id2*, *Nfactc1*, *Pdcd1*, *Lag3, Havcr2, Entpd1, Gzmk, Ccl3,Ccl4 and Ccl5*) (**Figure 2C, D**). Interestingly, bulk TILs from PD1d/2^V^-ACT clustered mostly in C1 (naïve/central-memory), while bulk PD1d/33-ACT TILs clustered mostly in C2, characterized by effector-memory genes (*Gzmk*, *Gzma*) (**Figure 2C, D**), confirming the orthogonal effects of the two cytokines seen above. Strikingly, bulk PD1d/2^V^/33-ACT TILs departed from all above states and clustered mostly in C5 (**Figure 2 C, D**), distinguished by low levels of exhaustion genes and high levels of cytotoxicity genes. The latter included *Gzmb* and notably high levels of *Gzmc*, a granzyme implicated in enhanced cytolytic function of T cells upon persistent antigenic stimulation and primary alloimmune responses, providing a non-redundant backup when granzyme B fails (Johnson et al. 2003; Cai et al. 2009; Getachew et al. 2008).

**Figure 2.**
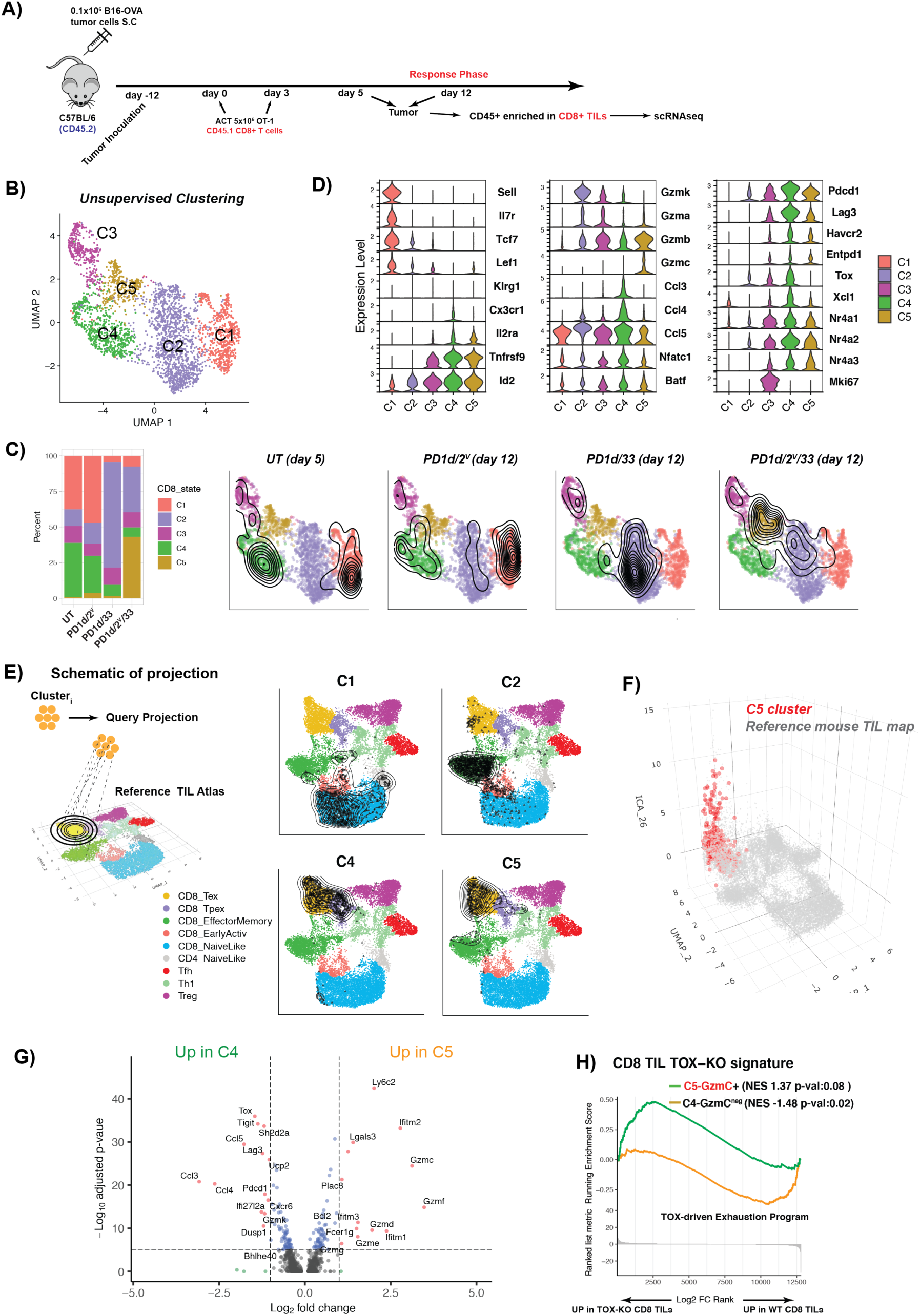
Orthogonal engineering induces a novel effector CD8^+^ T-cell state. **(A)** Experimental design. Briefly, mice with B16-OVA tumors were treated as indicated; then tumors were harvested on days 5 and 12 post ACT and a cell suspension of CD45^+^ enriched in CD8^+^TILs was obtained by FACS sorting and single cell sequenced using the 10X Genomics **(B-C)** Cluster composition per treatment and UMAP plot showing a low-dimensional representation of cell heterogeneity and unsupervised clustering results, where contour plots depict high cell density areas for each treatment. **(D)** Violin plots showing expression levels of important T cell markers **(E)** Projection of each cluster onto the reference TIL map using ProjecTILs. **(F)** ICA 26 effector component (Andreatta et al. 2021) for cells in the reference TIL atlas (gray) versus cluster C5 (red) samples, plotted on the z axis of the UMAP plot **(G)** Volcano Plot showing significant differentially expressed genes between clusters C4 and C5 **(H)** CD8 TIL Tox Knock-out (Scott et al 2019) Gene Signature Enrichment Analysis (GSEA) of Gzmc+C5 vs Gzmc^neg^ C4 cells.

Next, we sought to interpret these states based on prior knowledge; we applied ProjecTILs (Andreatta et al. 2021) (**Figure 2E**), “projecting” each cluster onto a reference atlas that comprises well-defined TIL states from untreated mouse tumors (Andreatta et al. 2021). Confirming the above state annotation using specific markers, the C1 cluster largely corresponded to the CD8^+^ naïve-like state, C2 to CD8^+^ effector memory state (EM), and C4 to canonical precursor (Pex) and terminal (Tex) exhausted states. Interestingly, C5 projected largely onto canonical Tex. Since this state was uniquely associated with tumor control, we investigated it further. Interrogation at a higher dimension using independent component analysis (ICA) revealed that C5 in fact deviated significantly from the reference Tex profile (**Figure 2F**), with a unique cytotoxic effector-like program (ICA26), that included several granzymes, notably GzmC (**Figure S3A**). Differential gene expression analysis confirmed that C5 is distinct from canonical exhaustion reported previously (McLane, Abdel-Hakeem, and Wherry 2019; Andreatta et al. 2021), or as seen here in C4 (**Figure 2G**). In addition to *Gzmc*, C5 TILs overexpressed several other granzymes as well as *Bcl2* and *Ly6c2* (**Figure 2G**), a marker of precursor CD8^+^ T cells which is absent in exhausted cells (Carmona et al. 2020). Relative to C4, C5 TILs also expressed more *Plac8*, implicated in optimal CD8^+^ T-cell response against viral infection (Slade et al. 2020). In addition, C5 TILs did not express *Cx3cr1* or *Klrg1,* distinctive markers of short-lived effector (SLEC) or transitory effector-like exhausted cells (Beltra et al. 2020; Chu and Zehn 2020; Hudson et al. 2019; Zander et al. 2019). Moreover, the inhibitory receptors *Pcdc1*, *Tigit* and *Lag3,* the exhaustion-associated chemokines *Ccl3*, *Ccl4* and *Ccl5*, as well as the exhaustion-associated transcription factor (TFs) *Bhle40, Nrp2a,Nfatc1,Nrp3a* (McLane, Abdel-Hakeem, and Wherry 2019; Chen, López-Moyado, et al. 2019) were lower in C5 compared to C4 cells (**Figure 2D, G**). Finally, *Tox,* a critical TF for the generation and maintenance of exhausted CD8^+^ T cells (Scott et al. 2019; Khan et al. 2019; Alfei et al. 2019; Yao et al. 2019; Seo et al. 2019) was nearly absent in C5 cells. Relative to C4, C5 cells were indeed enriched in the signature of *Tox*-knockout CD8^+^ TILs (Scott et al. 2019) (**Figure 2H**).

### C5 is a synthetic state uniquely acquired by orthogonally engineered CD8^+^ TILs

The above analyses were performed on total TILs harvested following ACT. We asked whether the novel C5 effector state was acquired both by transferred and by endogenous CD8^+^ TILs. We interrogated separately more FACS-sorted OT1 and endogenous CD44^+^CD8^+^ TILs at different time points following PD1d/2^V^/33-ACT (**Figure 3A**). C1, C2 and C5 distributed in an identical manner as before on ProjecTILs (**Figure S3B**), and C5 was again distinct from canonical Tex (**Figure S3B**). In addition, we could now distinguish the Pex from canonical Tex within C4 (**Figure 3B, S3C top part**). This analysis revealed two additional states, C6 and C7 (**Figure 3B**). C6 was characterized by a genomic profile enriched in Pex gene signature, while C7 was transcriptionally similar to *Pdcd1^low^* canonical memory cells recovered post-acute viral infection (**Figure S3C bottom part**).

**Figure 3.**
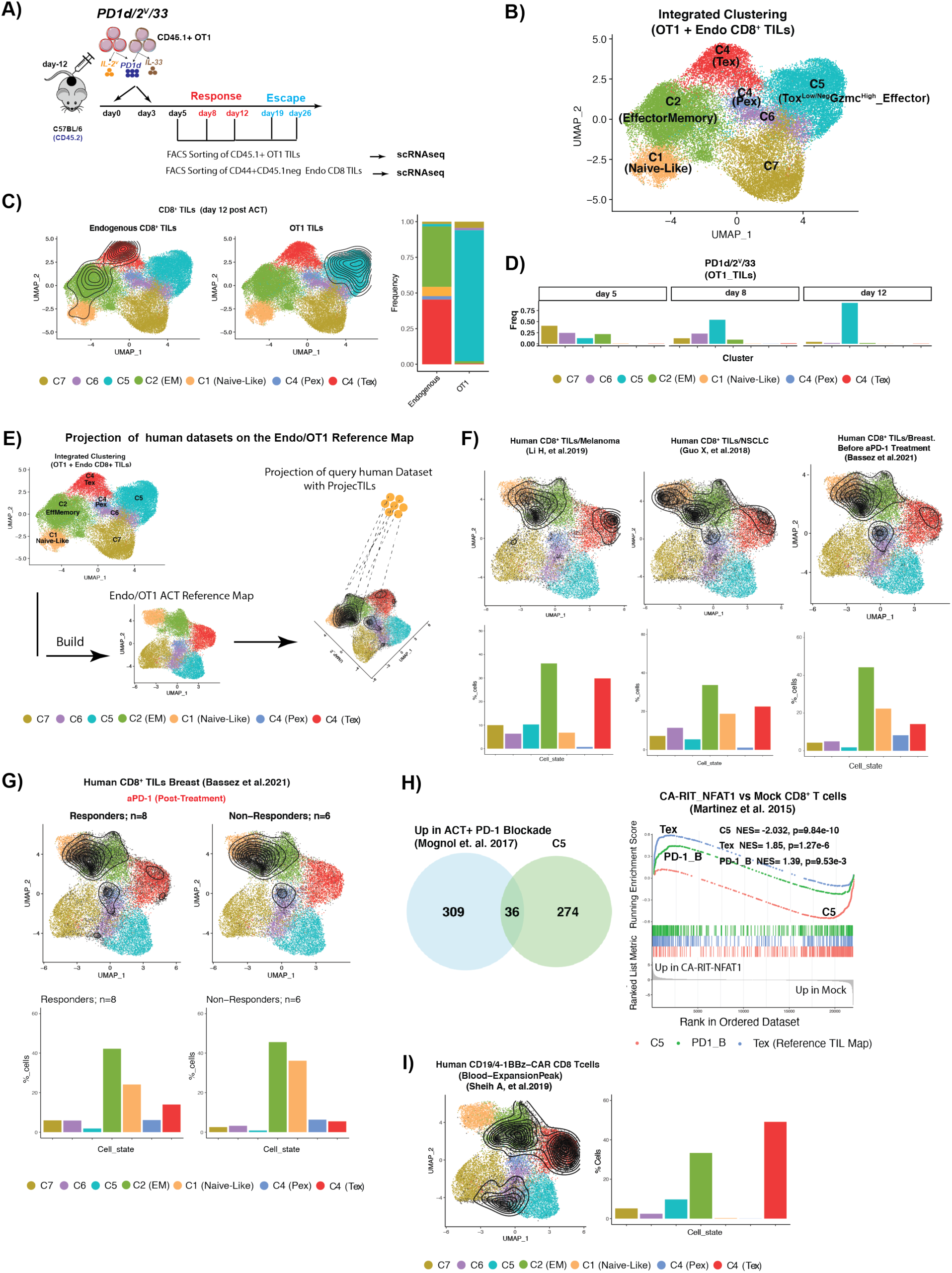
C5 is a novel synthetic state uniquely acquired by orthogonally engineered CD8^+^ TILs. **(A)** Experimental design. Briefly, mice with B16-OVA tumors were treated as indicated; then tumors were harvested on indicated time points and cell suspensions of either CD45.1^+^ OT1 and CD45.1^neg^ endogenous CD8 TILs were purified by FACS sorting and single cell sequenced using 10X Genomics **(B)** UMAP plot showing a low-dimensional representation cell heterogeneity and unsupervised clustering results. **(C)** UMAP plot showing the cluster composition of either Endogenous CD8 or OT1 TILs recovered 12 days post orthogonal ACT. Contour plots depict the clusters covered by each cell compartment. Bar plots are showing quantification of clusters covered by each cell compartment. **(D)** Cluster composition/time point of OT1 TILs recovered 5, 8 and 12 days post orthogonal ACT. **(E)** Schematic of projection of human CD8 TILs onto the OT1/Endogenous Transcriptomic space using ProjecTILs. (**F**) Projection of human CD8 TILs from three different types of tumors. Contour plots depict the clusters covered by each cell dataset and bar plots showed the quantification. (**G**) Projection of human CD8 TILs upon PD-1 blockade in responder versus non responder breast cancer patients onto the OT1/Endogenous Transcriptomic space using ProjecTILs. Contour plots depict the clusters covered by each cell dataset and bar plots are showing the cluster composition calculated by ProjecTILs. (**H**) Left: Venn diagram showing the overlap between genes upregulated in C5 (in the comparison showed in **Figure S3C**) and genes upregulated upon PD-1 blockade from Mognol et. al 2017. Right: CA-RIT-NFAT1 gene set enrichment analysis (GSEA) of canonical Tex TILs (genes downregulated in the comparison showed in **Figure S3B**), OT1 cells recovered upon PD-1 blockade (Mognol et. al. 2017.) and C5 cells (genes upregulated in the comparison showed in **Figure S3B**). This analysis shows that only Tex and OT1 cells recovered upon PD-1 blockade are enriched in the genes that characterize the severe exhaustion phenotype induced by the non-AP1 binding NFAT variant described in (Martinez et al. 2015). (**I**) Projection of human CAR T cells onto the OT1/Endogenous Transcriptomic space using ProjecTILs. Contour plots depict the clusters covered by each cell dataset and bar plots are showing the cluster composition calculated by ProjecTILs.

Importantly, only OT1 cells acquired the C5 effector state following PD1d/2^V^/33-ACT (**Figure 3C, S3D**). Indeed, by day 5 post ACT, most OT1 were distributed in C2 (EM), C6 (Pex-like) or C7 (canonical memory-like) states, and a minimal fraction had entered C5 (**Figure 3D**). By day 8, cells were mostly in C5 (54 %) and C6 (22%), and by day 12, >90% of cells were in C5, coinciding with tumor regression (**Figure 3D**). Conversely, endogenous CD44^+^CD8^+^ TILs mostly distributed across C4 (canonical Tex) and C2 (effector-memory) (**Figure 3B**, **S3D**). Thus, during tumor regression, adoptively transferred cells differentiate to the novel C5 effector-like state.

To find out whether this novel effector C5 state may arise during conventional immunotherapy, but has remained unnoticed, we used the ProjecTILs algorithm to place public domain scRNAseq TIL datasets in a new reference map that comprises the clusters identified above in endogenous CD8^+^ and OT1 TILs, shown in (**Figure 3**). Analysis of three different tumor type datasets showed that C5 is rare in human CD8^+^ TILs at baseline (i.e. no prior immunotherapy, **Figure 3F**). Furthermore, virtually no C5-like cluster was found in TILs from patients during response to anti-PD-1 (**Figure 3G**). Similarly, following adoptive transfer in combination with PD-1 blockade in mice (Mognol et al. 2017), OT1 TILs exhibited minimal overlap with C5, and were rather enriched in genes associated with the NFAT-dependent program of exhaustion (Martinez et al. 2015) (**Figure 3H**), consistent with the notion that PD-1 blockade does not reprogram TILs away from exhaustion (Held et al. 2019). Finally, we also interrogated human blood CD8^+^ CD19/4-1BBz CAR-T cells at peak expansion *in vivo* from four patients with durable CAR-T persistence following infusion (Sheih et al. 2020). We could not identify C5-like cells, and the majority of CAR-T cells corresponded to Tex-like cells (**Figure 3I**). Together, this data indicate that C5 is a synthetic novel state rarely seen following conventional immunotherapies but massively acquired *in vivo* by exogenous T cells following orthogonal-engineering ACT with PD1d, IL-2^V^ and IL-33.

### C5 OT1 TILs are polyfunctional effector cells with direct antitumor properties

We sought to characterize the cells ascribed to C5 effector state, which in our flow gating strategy we identified as PD-1^+^TOX^low/neg^GzmC^+^ cells. In untreated tumors, antigen-experienced (CD44^+^PD-1^+^) CD8^+^ TILs were largely TOX^+^ and GzmC^low^ (**Figure 4A**), consistent with canonical exhausted cells. During tumor regression post PD1d/2^V^/33-ACT, OT1 cells were mostly CD44^+^PD-1^+^ and highly enriched in TOX^low/neg^ GZMC^+^ cells (**Figure 4A**). Most GzmC^+^ were TCF1^neg^ (**Figure 4B**), expressed LY6C, but low/no KLRG1 or CX3CR1 (**Figure S3E)**, thus were altogether distinct from canonical Tex cells or SLEC.

**Figure 4.**
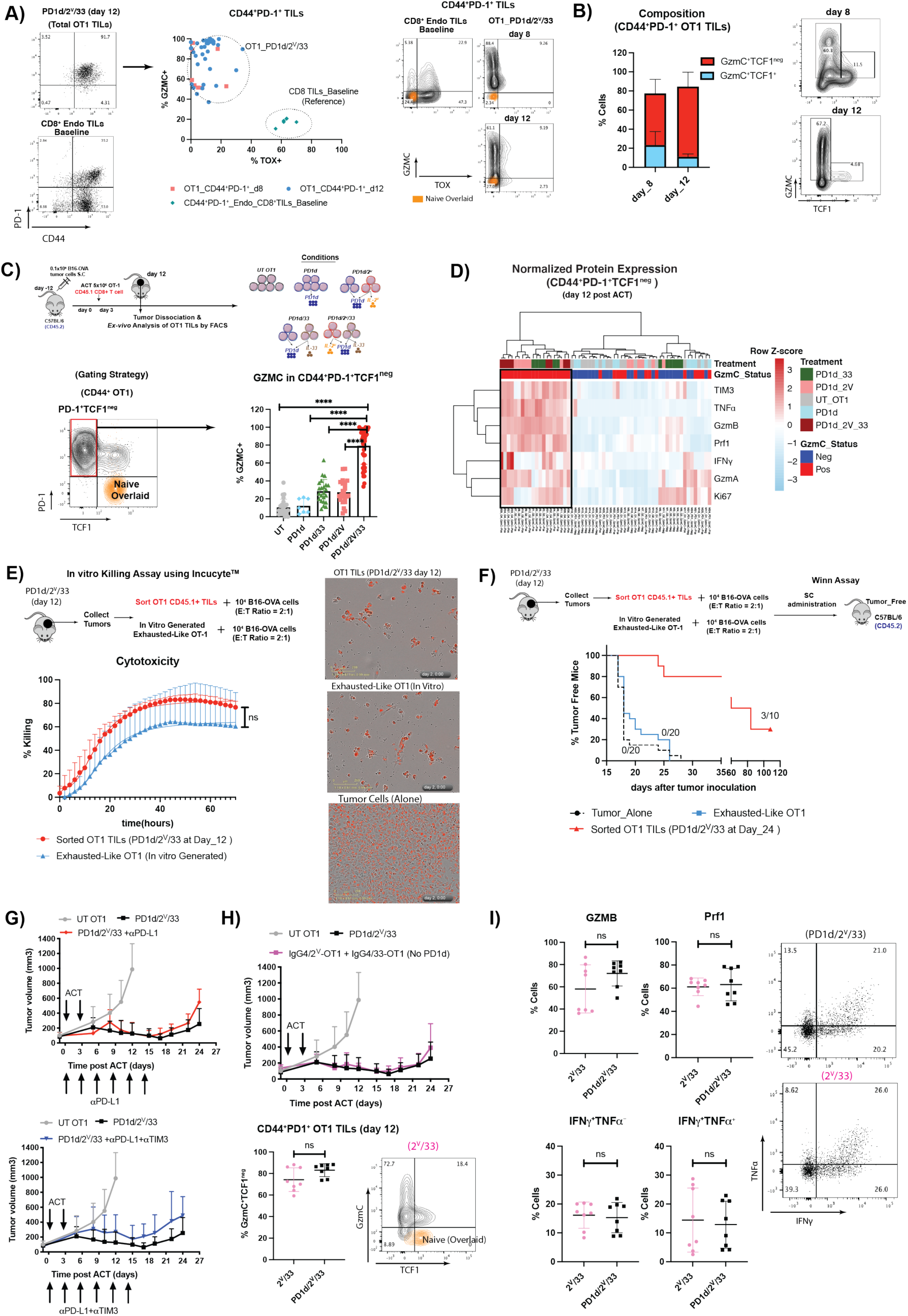
PD-1^+^TOX^Low/Neg^GzmC^+^ TCF1^neg^ OT1 TILs are polyfunctional effector cells with direct antitumor potential despite expression of coinhibitory receptors. **(A)** OT1 TILs from animals treated with orthogonal-engineered OT1 cells were analyzed at day 8 (two independent experiments n=4 mice/group) and day 12 (5 independent experiments n=5-6 mice/group) for the quantification of CD44, PD-1, TOX and GZMC by flow cytometry. CD8 TILs isolated from tumor-bearing mice at baseline were used as control (n=6 mice/group). On the right, a representative dot plot, naïve CD8 T cells isolated from the spleen of non-tumor bearing mice were used as negative control for the expression of the studied markers. **(B)** Left: GZMC and TCF1 expression in CD44^+^ PD-1^+^ TOX^Low/neg^ OT1 TILs recovered at days 8 and 12 post orthogonal-engineered OT1. Right: a representative contour plot. **(C)** Top: Experimental Design. Bottom: Analysis of GZMC expression in CD44^+^ PD-1^+^ TCF1^neg^ OT1 TILs recovered at day 12 post ACT from the different studied conditions (data from 5 independent experiments n=5-6 mice/group, except for PD1d, where we collected data from two independent experiments n=4 mice/exp). A Brown-Forsythe and Welch ANOVA test combined with Tukey Test to correct for multiple comparisons was used for comparing different groups * p<0.05, ** p<0.01, *** p<0.001, ****p<0.0001. **(D)** Heatmap showing the normalized (row scaled) protein expression of TIM3, TNFα, GZMB, PRF1, IFN,GZMA and KI67 in GZMC^+^ versus GZMC^neg^ CD44^+^ PD-1^+^ TCF1^neg^ OT1 TILs recovered at day 12. Each column represents an individual mouse (n=34 mice in total). (**E**) *In vitro* killing assay of sorted OT1 TILs 12 days post orthogonal ACT or *in vitro* generated exhausted OT-1 cells (as described in **Figure S3J**) against labelled B16-OVA tumors using Incucyte^TM^. **(F)** Top: Experimental design to evaluate the direct antitumor potential of CD44^+^ PD-1^+^ TOX^Low^/^Neg^ GZMC^+^ OT1 TILs recovered 12 post orthogonal ACT (data from two independent experiments, n=5 mice/exp for PD1d/2^V^/33 group and n=10 mice/experiment for the other two groups n=10 mice/experiment). Bottom Left: Percentage of tumor free mice after subcutaneous injection of tumor plus sorted OT1 TILs or in vitro generated exhausted OT-1. **(G)** Tumor growth control over time of B16.OVA tumor-bearing mice treated with PD1d/2^V^/33+ OT-1 cells in the presence or absence of 250*μ*g/mouse of antibodies specific for the indicated immune checkpoints administered i.p. beginning 1 day before 1^st^ cell transfer and maintained every three days maximum 6 doses **(H)** Top: Tumor growth control over time of B16.OVA tumor-bearing mice treated with PD1d/2^V^/33^+^ OT1 cells or with OT1 T cells gene-engineered for secreting only IL-2^V^ and IL-33 (no PD-1 ectodomain). Bottom: Percentage of GMZC^+^TCF1^neg^ effector-like cells in CD44^+^PD-1^+^ OT1 TILs recovered from mice treated with either PD1d/2^V^/33^+^ OT1 cells or IL-2^V^/IL-33 OT1 12 days post orthogonal ACT (2 independent experiments n=3-4 mice/group). **(I)** Expression of effector molecules in CD44^+^PD-1^+^GMZC^+^TCF1^neg^effector-like cells in OT1 TILs from **(H).** A two-tailed Student’s t test with Welch’s correction was used for comparing both groups in (**H** and **I**). ns: p>0.05. A Representative experiment out of two independent experiments (n= 6 animals/group) is shown for experiments in **F** and top **G**.

When we compared CD44^+^PD-1^+^TCF1^neg^ TILs on day 12 post ACT across all experimental conditions (**Figure 4C** **top**), GZMC^+^ cells were uniquely enriched in tumors following orthogonal ACT (**Figure 4C**), but were not detectable in the TDLN or spleens post ACT (**Figure S3F, G**), indicating that the C5 synthetic state develops specifically in tumors. Accordingly, co-administration of FTY720 did not affect the acquisition of this effector state in the tumors **(Figure S3H)**. Furthermore, unbiased analysis showed that PD-1^+^ GZMC^+^ TILs exhibited a unique phenotype of polyfunctional effector cells by FACS, with coexpression of GzmB, GzmA, perforin, TNFα, IFNγ and Ki67 despite also expressing TIM-3 (**Figure 4D** **and Figure S3I**), indicating that the synthetic new state C5, hallmarked by high expression of GzmC, is endowed with potent cytolytic and effector functions. To directly assess this, we harvested OT1 TILs from responding tumors following PD1d/2^V^/33-ACT, when 80%-90% of the OT1 are in C5, and tested their antitumor activity *in vitro* (**Figure 4E**) and *in vivo* using a Winn assay (**Figure 4F**). Unlike PD-1^+^ TOX^+^ OT1 control exhausted cells generated *in vitro* (Zhao et al. 2020) (**Figure S3J**), which exhibited cytolytic activity *in vitro*, but were unable to control tumor growth *in vivo*, OT1 TILs isolated from PD1d/2^V^/33-ACT tumors killed B16-OVA cells *in vitro* and led to tumor rejection *in vivo* (**Figure 4E, F**). Thus, OT1 effector TILs in C5 state are potent polyfunctional tumor-rejecting cells.

GZMC^+^ C5 OT1 cells expressed PD-1 and TIM-3 (**Figure 4A** **and Figure S3K**), yet they deviated from the canonical exhausted state. We sought to understand whether PD-1 or TIM3 expression translated to functional restriction. We thus combined PD1d/2^V^/33-ACT with therapeutic αPD-L1 or double αPD-L1/αTIM3 therapeutic antibody. Immune checkpoint blockade did not improve tumor control by orthogonal ACT (**Figure 4G**), indicating that PD-1 or TIM-3 inhibition were not important in the context of combinatorial IL-2^V^/IL-33 ACT. To confirm the lack of contribution of PD-1 blockade, we removed the PD-1 ectodomain from the PD-1_IgG4 decoy. Loss of PD1d did neither impact tumor control by IL-2^V^/IL-33-transduced OT1 cells nor impair the accumulation of CD44^+^ PD-1^+^GZMC^+^TCF1^neg^ OT1 TILs (**Figure 4H**). Indeed, OT1 TILs from 2^V^/33-ACT exhibited similar polyfunctional effector phenotype to TILs from PD1d/2^V^/33-ACT (**Figure 4I**). Collectively, these findings indicate that orthogonal engineering of T cells with a βγ-binding IL-2 and IL-33 in the immunocompetent host enables adoptively transferred CD8^+^ T cells to enter a novel synthetic effector state, which endows them with the ability to control tumors, and in which coinhibitory receptors – ordinarily associated with exhaustion – are expressed but remain functionally inconsequential.

### Orthogonal engineering ACT mobilizes endogenous CD8^+^ T cells against tumors

Given that mice were not lymphodepleted prior to ACT, we interrogated the endogenous tumor-infiltrating lymphocyte compartment. Strikingly, approximately 50% or more of total CD8^+^ TILs two weeks post-ACT were endogenous (CD45.1^neg^ CD45.2^+^, **Figure 5A**, see also **S2B, C**). These were particularly prominent in tumors treated with triple-engineered cells, where endogenous CD8^+^ T cells expanded preferentially over total CD4^+^ (**Figure S4A**), and the CD8^+^/CD4^+^FoxP3^+^ Treg cell ratio was the highest (**Figure 5B**). Antibody-mediated depletion of CD4^+^ T cells prior to PD1d/2^V^/33-ACT did not compromise, but instead significantly improved tumor control and mouse survival, indicating that endogenous CD4^+^ cells did not contribute to tumor control (**Figure 5C**, **Figure S4B)**. Lastly, given that recombinant IL-2^V^ mobilizes NK cells (Carmenate et al. 2013), we asked whether triple-engineered ACT activated NK-mediated antitumor immunity. NK-cell depletion did not impact tumor control, although tumor-infiltrating NK cells did increase post-ACT containing IL-2^V^, especially with PD1d/2^V^/33-ACT, but failed to display an activation phenotype (**Figure S4C-D).**

**Figure 5.**
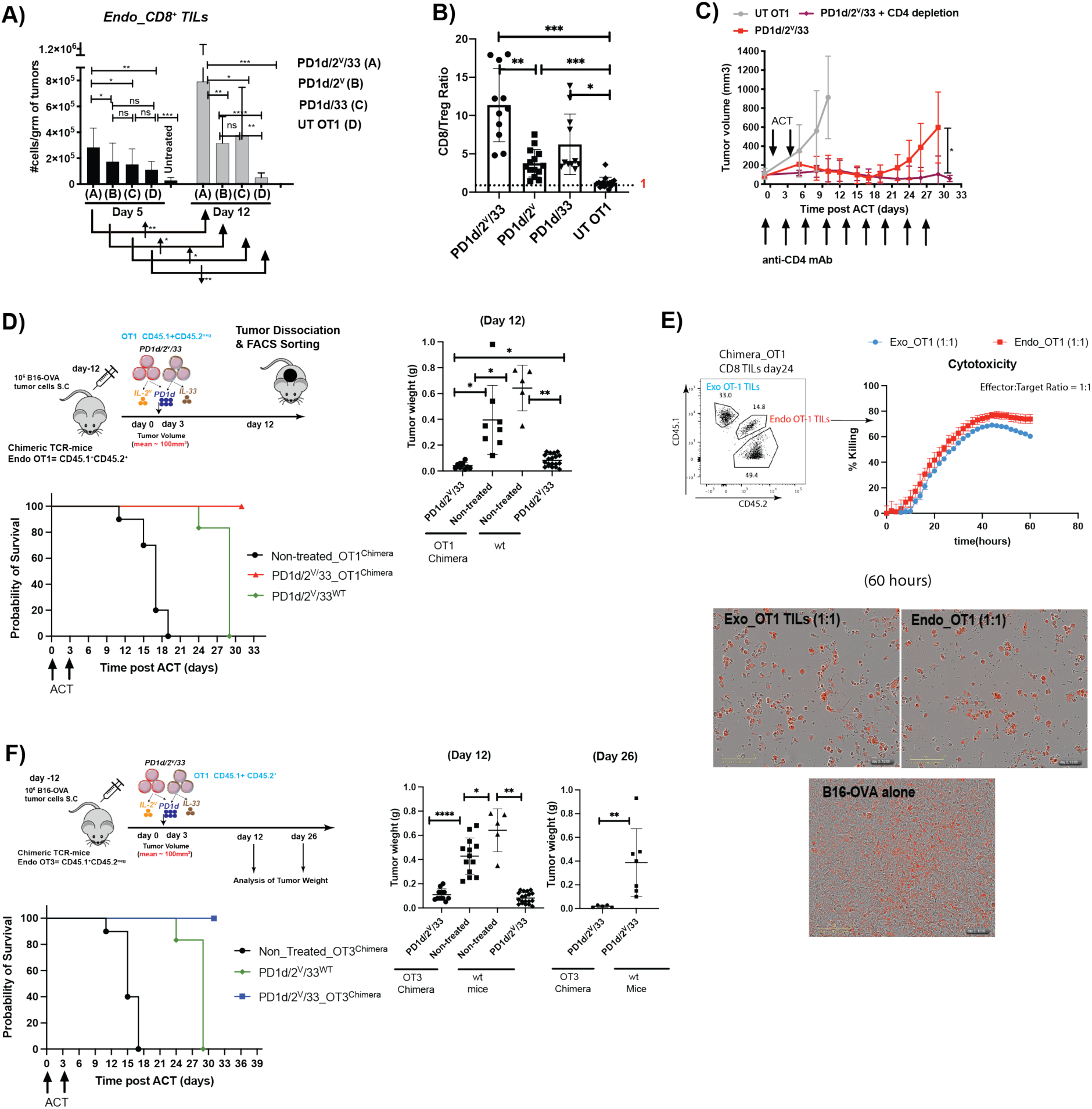
Orthogonal engineering ACT mobilizes endogenous CD8^+^ T cells against tumors. **(A)** Total number of endogenous CD45.1^neg^ CD8 TILs on days 5 and 12 post ACT. Mice with B16.OVA tumors were treated with either engineered or untransduced OT1 cells as indicated; then tumors were harvested on days 5 and 12 and cell quantification was performed by flow cytometry. Data are from three independent experiments (n >= 5 animals/group). **(B)** Mice with B16.OVA tumors were treated as indicated then tumors were harvested on day 12 post ACT, and Treg quantification was performed by flow cytometry. Data are from three independent experiments (n >= 5 animals/group). Shown are bar plots for CD8^+^/Treg ratio. **(C)** Tumor growth control over time of B16.OVA tumor-bearing mice treated with PD1d/2^V^/33+ OT-1 cells in the presence or absence of 250*μ*g/mouse of an αCD4 depleting antibody administered i.p. beginning 1 day before 1^st^ cell transfer and maintained every three days until day 33 post ACT. **(D)** Left: Experimental design of orthogonal ACT in OT1 chimeric mice. Right : weight of tumors harvested at day 12 post ACT from either chimeric OT1 mice or wt B16 mice treated in parallel with orthogonal ACT (two independent experiments, n=5-6 mice/group). Bottom: Survival curves. Tumor growth comparison was done using a Mann-Whitney Test **(E)** *In vitro* killing assay of FACS sorted exogenous and endogenous OT1 TILs against B16-OVA tumors using Incucyte^TM^. These cells were isolated from tumor-bearing OT1 chimeric mice 12 days post orthogonal ACT. Briefly, chimeric OT1 mice were generated as explained in materials and methods and two months after bone marrow transplantation were inoculated with B16.OVA tumors and treated with orthogonal engineered OT1 cells; then tumors were harvested on day 12 post ACT either exogenous (CD45.1^+^ CD45.2^neg^) or endogenous (CD45.1^+^CD45.2^+^) OT1 TILs were purified using FACS sorting. **(F)** Top: Experimental design of orthogonal ACT in OT3 chimeric mice. Bottom : weight of tumors harvested at days 12 and 26 post ACT from either chimeric OT3 mice or wt B16 mice treated or not in parallel with orthogonal ACT (two independent experiments, n=5-6 mice/group). Bottom: Survival curves. A Brown-Forsythe and Welch ANOVA test combined with Tukey Test to correct for multiple comparisons was used for comparing different groups in (A,B, D, F and G) and tumor volumes in (C). * p<0.05, ** p<0.01, *** p<0.001, ****p<0.0001.

We next asked whether endogenous CD8^+^ T cells contributed to tumor control after orthogonal ACT. Endogenous CD44^+^PD1^+^ CD8^+^ TILs harvested from responding tumors post PD1d/2^V^/33-ACT demonstrated cytolytic activity against B16-OVA cells *ex vivo* (Figure S4E), indicating that endogenous CD8^+^ TILs were also engaged against tumor. To further test the contribution of endogenous CD8^+^ TILs to tumor rejection upon orthogonal ACT, we used CD8 knockout mice, but these failed to engraft transferred OT1 cells (data not shown), providing a poor model for addressing the question. We thus resorted to mouse models in which endogenous T cells were either devoid of or enriched in tumor-specific precursors. In the former setting, we used transgenic TCR P14 tumor-bearing mice, where the bulk of endogenous CD8^+^ T cells are specific for a single lymphocytic choriomeningitis virus (LCMV) glycoprotein epitope gp_33–41_ and thus practically lack precursors directed against B16-OVA tumors. ACT with PD1d/2^V^/33-OT1 cells in P14 mice was indeed less effective than in wild-type mice (Figure S4F), suggesting some contribution of endogenous CD8^+^ T cells. In the opposite setting, irradiated CD45.1^neg^CD45.2^+^ mice were used to generate mixed wild-type/congenic OT1 TCR transgenic bone marrow chimeras (Figure S4G). Sixty days after bone marrow injection, when OT1 T cells represented around 30% of total CD8^+^ T cells in the blood, the reconstituted mice were inoculated with B16-OVA tumors. Twelve days post tumor inoculation, ∼20% of total CD8^+^ TILs were CD45.1^+^CD45.2^+^ OT1, which exhibited Pex or Tex features (Figure S4H). Orthogonal ACT with congenic PD1d/2^V^/33-transduced CD45.1^+^CD45.2^neg^ OT1 cells was more effective in OT1-chimeric mice than in wild-type recipient mice (**Figure 5D**). Furthermore, sorted endogenous (CD45.1^+^CD45.2^+^) OT1 TILs were as effective as exogenous OT1 (CD45.1^+^CD45.2^neg^) TILs in killing B16-OVA tumor cells *ex vivo* (**Figure 5E**). To further test whether low-affinity endogenous T cells also participate in tumor rejection upon orthogonal ACT, we repeated the same experiments using CD45.1^+^CD45.2^neg^ bone marrow to generate endogenous OT3 cells, which recognize the same OVA peptide SIINFEKL with lower affinity (Figure S4G). Orthogonal ACT with PD1d/2^V^/33 CD45.1^+^CD45.2^+^ congenic OT1 was more effective in inducing long-term tumor control in these OT3-chimeric mice than in wild-type recipient mice (**Figure 5F**). Thus, endogenous CD8^+^ T cells of high and low affinity are mobilized by orthogonal ACT and contribute to the antitumor response.

### Orthogonal engineering ACT reprograms endogenous CD8^+^ TILs to new states away from PD-1^+^TOX^+^ canonical exhaustion

At baseline as well as following orthogonal ACT in the wild-type host, endogenous C4 Tex-like cells featured a high proportion of clonally expanded cells (**Figure S5A**), indicating engagement against tumor. To learn whether these endogenous cells also deviated from canonical exhausted states under the influence of orthogonal ACT, we interrogated them longitudinally. Analysis of the exhausted compartment across many time points revealed four distinct states, one ascribable to TCF1^+^ Pex and the other three to Tex (**Figure 6A, top part**). At baseline and upon tumor escape (from day 19), endogenous CD8^+^ TILs were mostly in Tex1, characterized by *Tox, Nr4a2, Tigit, Gzmk, Pdcd1*, *Ccl3* and *Ccl4* (**Figure 6A**), consistent with the canonical *Tox*^high^*Nr4a2*^high^ exhaustion program (Scott et al. 2019; Chen, López-Moyado, et al. 2019). Following orthogonal ACT, endogenous Tex TILs departed transcriptionally from this state and acquired Tex2 (days 5 and 8) and later Tex3 (day 12). Neither subset upregulated *Cx3cr1* or *Tbx21* (**Figure 6A**), distinguishing them from transitory effector-like exhausted cells (Beltra et al. 2020). Tex2 and Tex3 exhibited significant enrichment in C5-related genes relative to Tex1 (**Figure 6B**). Relative to Tex1, Tex2 downregulated canonical Tex TFs *Tox* and *Nr4a2* and Tex markers *Gzmk, Ccl3, Ccl4* and *Ccl5* (**Figure 6C**) while Tex3 TILs also upregulated *Gzmb*, *Prf1*, *Plac8, Serpinb9*, *Tnfrsf9* (4-1BB) and *Ly6c2* (**Figure S5B**), suggesting better cytolytic function, also seen by pathway analysis (**Figure S5C**). Thus, under the influence of orthogonal ACT, endogenous CD8^+^ TILs acquired an “infectious” activated-Tex state, transcriptionally closer to C5, with low *Tox* and upregulated effector machinery. Importantly, even though highly expanded clonotypes could distribute across more than one Tex subtype (**Figure S5D**), they mostly acquired Tex2 and Tex3 during tumor regression.

**Figure 6.**
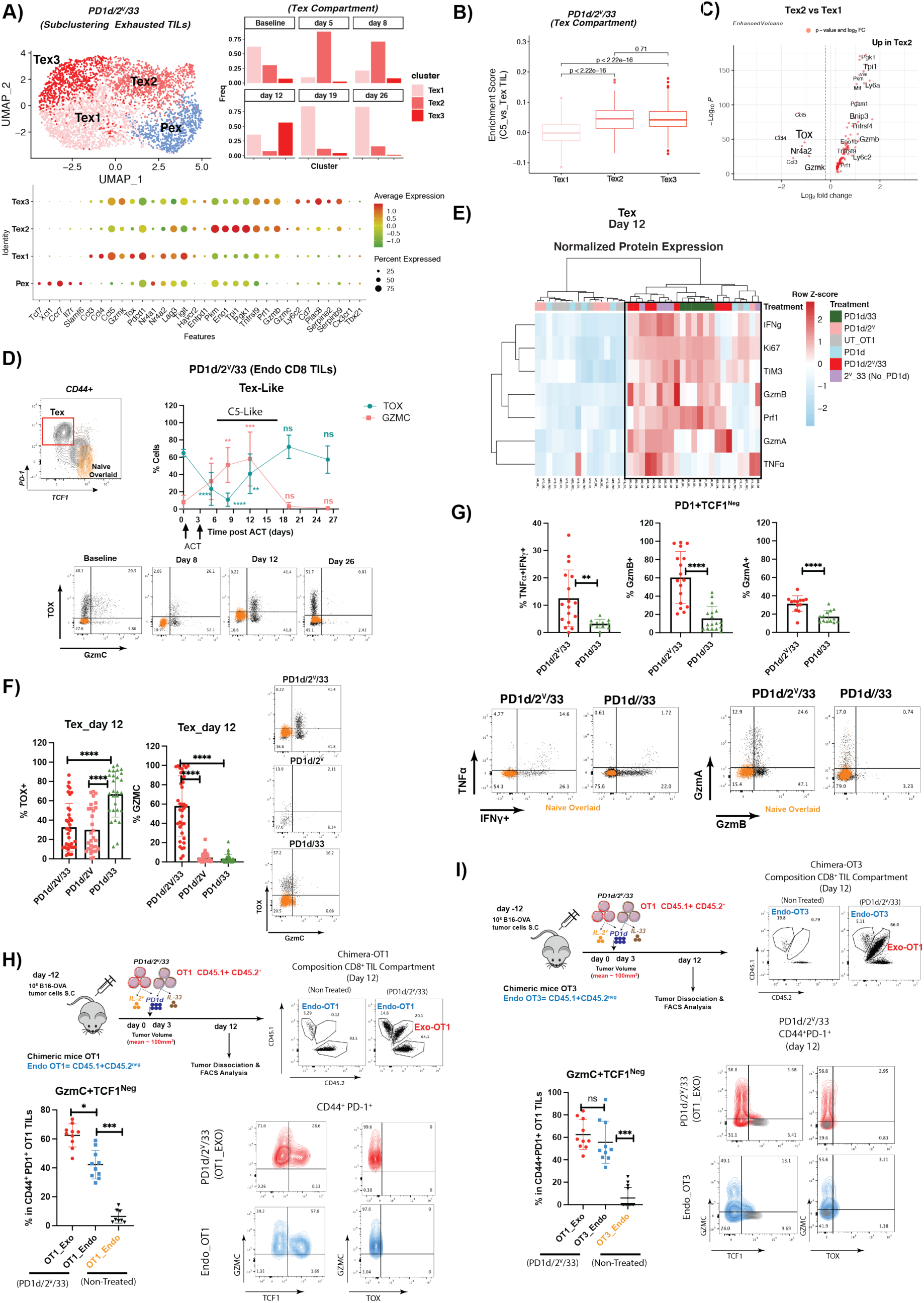
Orthogonal engineering reprograms endogenous Tex TILs. **(A)** Top Left: UMAP plot showing a low-dimensional representation of cell heterogeneity and unsupervised clustering results of endogenous exhausted CD8 TILs (Pex and Tex) expanded by orthogonal ACT across all studied time points (from baseline to day 26 post ACT). Top Right: cluster composition/time point. Bottom part: dot plot showing clusters-specific markers. (**B**) Enrichment score of C5 gene signature (genes upregulated in the comparison showed in **Figure S3C)** in endogenous Tex subtypes (the enrichment score was calculated with AUCell). A non-parametric Kruskal Wallis test was used for multiple comparisons * p<0.05, ** p<0.01, *** p<0.001, ****p<0.0001. **(C)** Volcano plot showing differentially expressed genes between Tex2 and Tex1 subtypes. (**D**) Expression TOX and GZMC in endogenous Tex cells harvested at different time points following orthogonal ACT and defined as depicted in the dot plot based on the expression of TCF1 and PD-1. A two-tailed Student’s t test with Welch’s correction was used for comparing expression of each marker, in each time point relative to baseline Tex TILs * p<0.05, ** p<0.01, *** p<0.001, ****p<0.0001. Naïve CD8 T cells isolated from the spleen of non-tumor bearing mice were used as negative control for the expression of the studied markers. **(E)** Heatmap showing the normalized (row scaled) expression of IFN, Ki67, TIM3, GZMB, PRF1, GZMA and TNFα in CD44^+^ PD-1^+^ TCF1^neg^ endogenous Tex TILs recovered at day 12 from each studied condition. Each column represents an individual mouse (n=34 mice in total). **(F)** Analysis of TOX and GZMC expression in endogenous Tex TILs recovered at day 12 post ACT from mice treated with PD1d/2^V^/33, PD1d/2^V^ and PD1d/33. Data collected from 5 independent experiments n= 5-6 mice/group. One-way ANOVA test in combination with a Dunnet Test to correct for multiple comparisons was used * p<0.05, ** p<0.01, *** p<0.001, ****p<0.0001. **(G)** Comparison of endogenous Tex CD8^+^ TILs harvested at day 12 post ACT from PD1d/2^V^/33 or PD1d/33-treated mice in terms of expression of effector molecules. Data collected from 3 independent experiments n= 3-5 mice/group. A two-tailed Student’s t test with Welch’s correction was used for comparing expression of each marker, in each time point relative to baseline Tex TILs * p<0.05, ** p<0.01, *** p<0.001, ****p<0.0001. Naïve CD8 T cells isolated from the spleen of non-tumor bearing mice were used as negative control for the expression of the studied markers. **(H-I)** Analysis of TOX, TCF1 and GZMC expression in CD44^+^ PD-1^+^ endogenous OT1 **(H)** and OT3 **(I)** TILs isolated from chimeric mice (two independent experiments n= 5-6 mice/group. A Brown-Forsythe and Welch ANOVA test combined with Tukey Test to correct for multiple comparisons was used for comparing different groups. * p<0.05, ** p<0.01, *** p<0.001, ****p<0.0001

Next, we confirmed by FACS that endogenous Tex-like CD44^+^PD-1^+^TCF1^neg^ TILs indeed downregulated TOX following orthogonal ACT (**Figure 6D**). Although we could not detect *Gzmc* mRNA in most Tex2 or Tex3 cells by scRNAseq, endogenous CD44^+^PD-1^+^TCF1^neg^ TILs from day 5 to day 12 (peak), unlike baseline Tex TILs, expressed GZMC protein (**Figure 6D**), albeit at lower levels than OT1 cells from the same responding tumors (**Figure S5E**). Endogenous PD-1^+^GZMC^+^ TOX^low^ CD8^+^ TILs cells found post orthogonal ACT co-expressed TNFα and IFNγ, as well as GzmA, GzmB and perforin (**Figure 6E**), indicating that they were superior effector cells than baseline endogenous Tex TILs, consistent with partaking in tumor attack (**Figure S5F**).

To understand the contribution of the individual engineering components in imparting such state transitions to endogenous CD8^+^ TILs, we compared Tex-like cells from different ACT conditions. Importantly, downregulation of TOX in endogenous CD8^+^ Tex cells was seen with any ACT that included IL-2^V^-OT1 cells (**Figure 6F** **left**), suggesting that IL-2^V^ drove suppression of TOX in endogenous TILs. However, GZMC, the other hallmark of the new state, was upregulated by endogenous CD8^+^ TILs only following orthogonal ACT (**Figure 6F** **right**), indicating that the synergistic interaction between IL-2^V^ and IL-33 (but not PD1d (**Figure S5E**)) was required. Furthermore, the endogenous Tex-like cells isolated post IL33-ACT upregulated multiple effector molecules, but not GZMC (**Figure 6E, F**), and these were highest post orthogonal ACT (**Figure 6G**). Elimination of PD1d did not impair the acquisition of these effector functions (**Figure 6E**), confirming that PD-1 blockade is not required for the functional reinvigoration of endogenous Tex TIL by orthogonal ACT. Importantly, we did not detect TCF1^neg^ endogenous C5 effector-like cells in the TDLN during the response phase (**Figure S5G**), and accordingly, co-administration of FTY720 did not alter the fraction of endogenous CD44^+^PD1^+^TOX^neg/low^ GzmC^+^TCF1^neg^ Tex TILs **(Figure S5H**), suggesting that Tex reprograming is specifically induced locally in the TME.

Natural endogenous tumor-specific T cells are expected to harbor TCRs across a range of affinities. We found that orthogonal engineering mobilizes high-as well as low-affinity tumor-specific endogenous CD8^+^ TILs and reprograms both to a similar C5-like effector state. Indeed, interrogating chimeric mice engrafted as above with either OT1 or OT3 cells, in which endogenous CD8^+^ TILs recognize the tumor peptide SIINFEKL with high or low affinity respectively, we saw that a large fraction of endogenous cells, OT1 or OT3, acquired a PD1^+^TCF1^neg^GZMC^+^TOX^low/neg^ C5-like phenotype (**Figure 6H, I**). Interestingly, the baseline state of these cells was different, as the high-affinity OT1 TILs were mostly TOX^+^ canonical exhausted, while low-affinity OT3 TILs were mostly CD44^+^PD-1^neg^TOX^neg^TCF1^+^ precursor-like cells (**Figure S5I**). However, under the influence of orthogonal ACT, both cells reprogrammed to C5-like effector cells.

### Orthogonally engineered ACT reprograms PD-1^+^ TCF1^+^ precursor-like TILs within the tumor microenvironment

As detailed above, the accumulation of TCF1^+^ OT1 or endogenous CD8^+^ precursors in tumors and subsequent generation of TCF1^neg^ effector CD8^+^ TILs was independent of lymphoid organs. Furthermore, transferred OT1 cells acquired specifically the novel C5 effector state in the TME. Trajectory and pseudotime inference analysis of OT1 cells suggests that they likely transitioned from the original naïve/central memory-like (C1) and effector-memory (C2) states of transferred cells to C5 through an intermediate C6 precursor-like state, bypassing the canonical Pex state observed in endogenous CD8^+^ TILs from baseline tumors (**Figure S6A**). The C6 state expressed *Pdcd1* together with the stemness-related markers *Ccr7, Tcf7* and *Il7r* (**Figure S6B, C**), but was different from canonical *Pdcd1^high^* Pex TILs. Indeed, relative to the latter, C6 TILs exhibited significant downregulation of the exhaustion-associated transcription factor *Tox* and, like C5, overexpressed *Gzmc, Gzmb, Ly6c2, Ctla2a* and *Ifitm2* (**Figure S6D**). Accordingly, most PD-1^+^TCF1^+^ OT1 TILs during the tumor control phase post PD1d/2^V^/33-ACT expressed higher GZMC, but lower TOX in comparison with canonical TCF1^+^PD-1^+^ Pex-like cells from baseline tumors (**Figure S6E**). Remarkably, only PD-1^+^TCF1^+^ OT1 TILs generated upon orthogonal ACT expressed significant levels of GZMC in comparison with the similar population recovered from PD1d/2^V^ or PD1d/33-treated mice at day 12 (**Figure S6F**). Thus, similar to cells committed to terminal exhaustion which is preceded by a PD-1^+^ precursor Pex state (Siddiqui et al. 2019; Miller et al. 2019), the alternate C5-synthetic effector differentiation program achieved by orthogonal engineering cells is also committed early, and transitions through a PD-1^+^TCF1^+^TOX^low/neg^ precursor/stem-like state, harboring a transcriptional program that is not consistent with canonical Pex.

Notably, during the response phase upon orthogonal engineering the endogenous CD8^+^ Pex TILs also acquired a state resembling C6 (**Figure S6G**). Like C6, these cells downregulated *Tox, Nr4a2, Xcl1* and upregulated *Ly6c2* (**Figure S6H**). Furthermore, following orthogonal engineering we also observed expression of GzmC by FACS in endogenous Pex CD8^+^ TILs (**Figure S6I**), which also downregulated TOX, most significant on day 8 post ACT (**Figure S6J**). Similarly to the reprogramming seen in endogenous CD8^+^ Tex above, we found that downregulation of TOX in endogenous TCF1^+^PD-1^+^CD8^+^ Pex cells was observed only in ACT conditions that included IL-2^V^-transduced cells, while expression of GzmC required concomitant IL-33. Thus, orthogonal ACT also reprograms endogenous Pex-like CD8^+^ TILs, deviating them in a coordinated manner away from canonical TOX^+^ PD-1^+^ Pex state.

### Tumor escape to orthogonal T-cell engineering is associated with loss of synthetic conditions leading to C5 state

Given the marked efficacy of orthogonally engineered T cells to control tumors, we asked what could be the reasons for tumor escape. We assessed the evolution of both OT1 (**Figure 7A, B**) and endogenous CD8^+^ states (**Figure 7A, C**) longitudinally from response to progression following triple-engineered ACT. Remarkably, OT1 migrated away from the dominant C5 state (tumor control, day 12) to the *Ccr7*^high^*Pdcd1*^low^*Tox*^low^*Gzmc*^low^ C7 state **(Figure 7B, S7A)**. Similarly, endogenous TILs migrated away from being highly expanded C4 Tex (and mostly Tex3 on day 12, see **Figure 6A**) to predominantly effector-memory cells (**Figure 7C**), including the highly expanded clonotypes (**Figure 7D**). Thus, tumor escape was not associated with accumulation of dysfunctional Tex, but rather with a loss of differentiation to highly functional effector states of both engineered (C5) and endogenous cells (Tex2/Tex3). By flow cytometry we confirmed a marked loss of the characteristic CD44^+^PD-1^+^TOX^Low^/^neg^GzmC*^+^*OT1 TILs (corresponding to C5) upon tumor escape (**Figure 7E**), coinciding with a marked enrichment in TCF1^+^PD-1^neg/low^ CD8^+^ T cells in both OT1 and endogenous cells (**Figure 7F**), consistent with the acquisition of C7 and effector-memory states, respectively. Residual TCF1^neg^ effector cells during escape lost GzmC and their polyfunctional features relative to GZMC^+^TCF1^neg^ PD-1^+^CD8^+^ T cells detected during tumor control (the latter in **Figure S7B)**.

**Figure 7.**
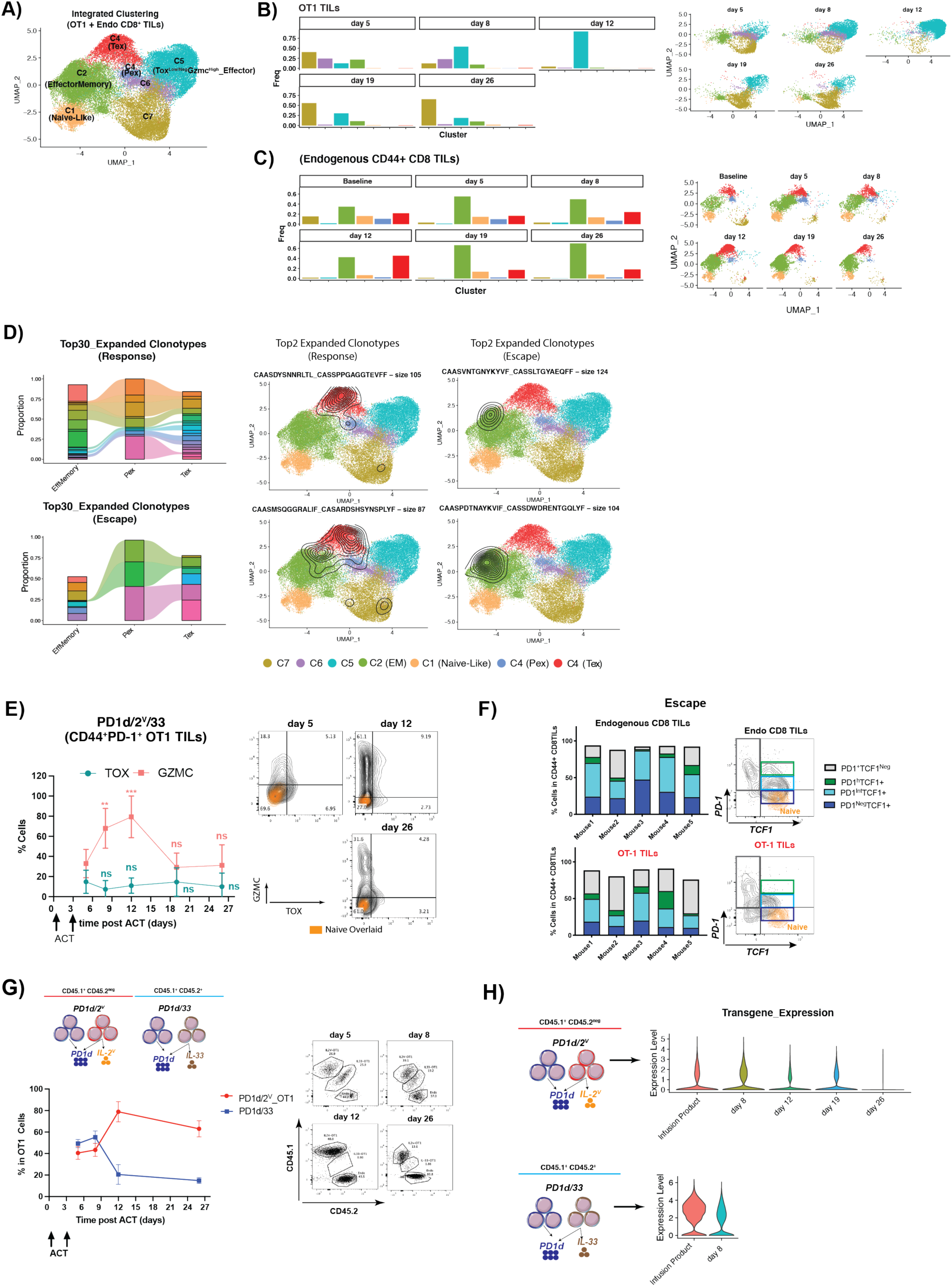
Tumor escape to orthogonal T-cell engineered ACT is associated with cessation of intratumoral differentiation to effector-like states in TILs. **(A)** UMAP plot showing a low-dimensional representation cell heterogeneity and unsupervised clustering results combined OT1 and endogenous CD8 TILs. **(B-C)** Cluster composition of TILs overtime in each compartment/time point. (**D**) Cell state distribution of the top 30 most expanded endogenous clonotypes during response or escape. On the right, a graphical example of the top 2 clonotypes/condition, where contour plots depict high cell density areas for each clonotype. (**E**) Longitudinal analysis of TOX and GZMC expression in exogenous CD44^+^PD-1^+^OT1 TILs harvested at different time points following orthogonal ACT. A two-tailed Student’s t test with Welch’s correction was used for comparing expression of each marker in each time point relative to OT1 harvested at day 5 post ACT * p<0.05, ** p<0.01, *** p<0.001, ****p<0.0001. Naïve CD8 T cells isolated from the spleen of non-tumor bearing mice were used as negative control for the expression of the studied markers. **(F)** Analysis of exogenous and endogenous CD8^+^ TILs harvested during escape (day 26) based on PD-1 and TCF1 expression. **(G)** Analysis of intratumoral persistence of congenic OT1 TILs transduced with either PD1d/2^V^ or PD1d/33. **(H)** Analysis of transgene expression overtime at the single cell level in OT1 TILs from different conditions. UT OT1 TILs were used as negative control for the expression.

Since TILs with predominant precursor-like states were characteristic of ACT with IL-2^V^-transduced cells lacking IL-33 coexpression, where PD1d/IL-2^V^ secreting OT1 cells were expanded but unable to transition to TCF1^neg^ effector-like cells (**Figure 1G**), we reasoned that the arrest in less differentiated states upon tumor progression might be a consequence of attrition of IL-33 secreting OT1 cells. To assess this hypothesis, we conducted orthogonal ACT using CD45.2^neg^ PD1d/2^V^-OT1 cells and congenic CD45.2^+^ PD1d/33-OT1 T cells (**Figure 7G**). Notably, the proportion of PD1d/33-secreting OT1 cells dropped dramatically by day 12 and beyond (**Figure 7G**), while the PD1d/2^V^-transduced OT1 TILs persisted. Thus, tumor escape could be attributed to the loss of the synthetic effector states in transferred (C5) and endogenous CD8^+^ TILs (Tex2/3), which in turn we attribute to the lack of persistence of IL-33-transduced engineered cells. We also found that in the remaining OT1 TILs, the expression of the transgene partially decreased after day 12 (**Figure 7H**), suggesting that the remaining IL-2^V^ in the TME was sufficient to ensure persistence of IL2^V^-secreting but not IL-33 secreting OT1 cells, leading residual IL2^V^-secreting OT1 TILs to arrest in the C7 state, which was the final state acquired by in the absence of IL-33 secretion (**Figure S7C, D**). Indeed, confirming the lack of paracrine effect of residual IL2^V^-engineered OT1 TILs, residual endogenous Pex (CD44^+^PD-1^+^TCF1^+^) and Tex (CD44^+^PD-1^+^TCF1^neg^) CD8^+^ TILs converted to canonical Pex and Tex1respectively, expressing high levels of TOX and no GZMC (see **Figure 6D** **and Figure S7E**).

## Discussion

We provide evidence that orthogonal combinatorial T-cell engineering overcomes homeostatic barriers to engraftment, reprograms TILs as well as the tumor microenvironment and mediates solid tumor regression. Adoptive T-cell transfer therapy of solid tumors in the immune competent host offers unique advantages as it may substantially reduce the toxicity and costs of present-time ACT, and may enable leveraging the host’s immune system. Indeed, we show that endogenous tumor specific CD8^+^ cells were also mobilized and engaged in tumor regression.

As a proof of principle of orthogonal T-cell engineering, here we successfully combined two main secreted components: an IL-2 variant binding to β*/*γ but not αIL-2R, together with IL-33. This combination brought about synthetic states, both in the adoptively transferred gene-engineered T cells as well as in endogenous CD8^+^ TILs. The acquired effector states in endogenous CD8^+^ TILs departed fundamentally from the previously known terminal exhausted state, the ubiquitous hallmark of tumor-engaged TIL in humans and mice. Notably, adoptively transferred gene-engineered CD8^+^ T cells acquired a unique novel non-canonical state, reproducibly associated with marked local CD8^+^ T-cell expansion in tumors, potent effector function, and tumor control. This state is characterized by expression of multiple granzymes (most prominently heralded by the unique granzyme-C), and significant downregulation of several transcription factors involved in CD8^+^ TIL dysfunction such as *Nfatc1*, *Nrp4a1* and *Nrp4a2* (Chen, López-Moyado, et al. 2019; Martinez et al. 2015). Notably, this effector state was also distinguished by expression of inhibitory receptors, but low or absent expression of TOX – a transcription factor that is critical for the generation and maintenance of exhausted CD8^+^ T-cell populations during chronic viral infection and in cancer (Mann and Kaech 2019; Scott et al. 2019; Alfei et al. 2019; Yao et al. 2019; Seo et al. 2019; Khan et al. 2019). These findings indicate that PD-1 and other co-inhibitory receptors can be upregulated in the context of sustained antigen stimulation independently of TOX, unlike previous reports describing TOX dependency for this in canonical exhausted CD8^+^ T cells (Mann and Kaech 2019; Scott et al. 2019; Alfei et al. 2019; Yao et al. 2019; Seo et al. 2019; Khan et al. 2019). Despite expressing PD-1 and TIM3, TILs in the synthetic C5 effector state were not constrained by these canonical inhibitory checkpoints; blockade of PD-1 and/or TIM3, or addition of a PD-1 decoy gene engineering module in the cells, was functionally inconsequential. Thus, the combination of IL-2^V^ and IL-33 drove TILs into a synthetic state of superior effector cells that were liberated of canonical exhaustion constraints.

As recently shown, the CD8^+^ T-cell exhaustion program is stably enforced in the PD-1^+^TCF1^+^ progenitor compartment by expression of TOX (Mann and Kaech 2019; Scott et al. 2019; Alfei et al. 2019; Yao et al. 2019; Seo et al. 2019; Khan et al. 2019; Kallies, Zehn, and Utzschneider 2020; Beltra et al. 2020; Chu and Zehn 2020). In the context of orthogonal engineering, adoptively transferred engineered CD8^+^ TILs also transitioned through a PD-1^+^TCF1^+^ precursor-like state. However, contrary to canonical Pex cells, transferred precursor cells exhibited low/no TOX and upregulated granzyme-C, a marker not detected hitherto in the canonical precursor exhausted compartment. Thus, the induced PD-1^+^TCF1^+^ progenitor-like C6 state also diverges from the canonical exhaustion fates described in the context of chronic viral infection and in cancer. Moreover, the novel synthetic effector state diverges also from the transitory effector-like exhausted state (CX3CR1^+^TIM3^+^PD-1^+^) (Beltra et al. 2020; Chu and Zehn 2020; Hudson et al. 2019; Zander et al. 2019), which arises from canonical Pex cells and does not upregulate GZMC (Beltra et al. 2020; Chu and Zehn 2020; Hudson et al. 2019; Zander et al. 2019), but expresses *Cx3cr1* and *Klrg1*, two markers not expressed by the TIM3^+^PD-1^+^GZMC^+^ synthetic effector state induced by orthogonal engineering. In addition, while CD4^+^ T-cell help is required for the formation of the CX3CR1^+^ effector state (Zander et al. 2019), these cells appeared to be dispensable or even deleterious in the context of our approach. In fact, query of public human and mouse TIL data revealed that the generated synthetic state has never been reported to date. Taken together, we hypothesize that both TCF1^+^ and TCF1^neg^ PD-1^+^GzmC^+^CD8^+^ TILs represent the precursor and effector states, respectively, of a novel PD-1^+^TOX-indifferent CD8^+^ T-cell synthetic differentiation program.

Strikingly, the expression of IL-2^V^ and IL-33 by engineered T cells reprogrammed endogenous CD8^+^ TILs, which also departed from the canonical TOX^+^ exhaustion program and developed transcriptomic cell states similar to those acquired by the engineered TILs. Such reprograming was seen mostly in clonally expanded TILs and was notably characterized by downregulation of TOX, and concomitant upregulation of GZMC along with genes encoding multiple cytotoxicity-related molecules. Remarkably, tumor-specific endogenous TILs with both high-affinity as well as low-affinity TCRs converted to this PD-1^+^TOX^low/neg^ GZMC^+^ C5-like superior effector state, and contributed to tumor regression. This is in contrast with the report that subdominant antigen specific CD8^+^ TILs do not preferentially benefit from anti-PD-1/CTLA4 therapy (Burger et al. 2021). Thus, convergence of both endogenous and exogenous CD8 TILs into similar effector-like states during tumor regression is a key property of orthogonal ACT.

The therapeutic manipulation of TOX has been proposed as a promising strategy to abrogate T-cell exhaustion in the context of cancer (Mann and Kaech 2019). This notion is supported by the improved functionality of CAR-T cells upon TOX knockdown or deletion of TOX2 (Seo et al. 2019), and the strengthening of anti-tumor T-cell responses reported in heterozygous deletion of TOX (Khan et al. 2019). In this study we show a novel approach to reach synthetic cell states in which TOX is suppressed transcriptionally without genome editing or knockdown, in both adoptively transferred as well as endogenous CD8^+^ T cells. Careful analysis of the states reached under different experimental conditions revealed that IL-2^V^ was primarily responsible for promoting CD8^+^ T-cell stemness and was associated with downregulation of TOX in PD-1^+^ exhausted cells. This effect was observed under all *in vivo* conditions including cells transduced with the IL-2^V^ module, and extended also to exogenous OT1 cells that were not transduced with IL-2^V^ as well as to endogenous TILs. Furthermore the effect of IL-2^V^ was already observed *in vitro* during T-cell expansion. How IL-2 signaling directed through the IL-2Rβγ receptor chains achieve suppression of TOX will require additional investigation, however it appears to be part of a transcriptional and/or epigenetic reprogramming that constituted the base for producing a novel effector state that profoundly deviates from canonical exhaustion upon antigen engagement. Finally, the fact that IL-2^V^ is associated with high level of persistence of TOX^low/neg^ PD-1^+^ CD8^+^ T cells in the context of sustained antigen stimulation is also a novel and remarkable finding, since this property is strictly depend on TCR-induced TOX expression in canonical exhausted CD8^+^ T cells (Alfei et al. 2019; Scott et al. 2019; Khan et al. 2019; Seo et al. 2019) . This suggests that IL-2 signaling directed through the IL-2Rβγ receptor chains could also play a key role in making CD8^+^ T cells less dependent on sustained TCR stimulation to survive in contexts of chronic antigen exposure.

The simultaneous expression of IL-33 *in vivo* was required to produce the novel synthetic effector state associated with superior antitumor functions. In its absence, cells exposed to IL-2^V^ remained in a state resembling memory endowed with stemness. IL-33 alone in the absence of IL-2^V^ led to activation of endogenous Tex-like cells, but this was associated with short persistence capability of transferred IL-33-secreting OT1. When combined with IL-2^V^ *in vivo*, IL-33 enabled GzmC upregulation and drove Tcf1 suppression, allowing OT1 cells to exit the precursor state and differentiate to polyfunctional PD-1^+^TOX^low/neg^ effector cells. However, such cells never entered to the canonical TOX^+^ exhaustion program (Alfei et al. 2019; Scott et al. 2019), apparently thanks to the effects of long-term exposure (*in vitro* and *in vivo*) to IL-2^V^. We speculate that the latter reprograms T cells enabling them to avoid the canonical TOX^+^ exhaustion program upon sustained antigen stimulation. However, our data also confirms that TOX downregulation in CD8^+^ T cells is not enough to effectively control tumors (Scott et al. 2019), since TILs recovered from mice treated with PD1d/IL2^V^ were also TOX ^low/neg^ but this group did not achieve significant tumor control. Thus, how IL-33 achieves further reprogramming of TILs toward a polyfunctional TOX^neg^ PD-1^+^ TCF1^neg^ effector state with direct antitumor potential remains to be elucidated. Activation of antigen-presenting cells in addition to inflammatory cues are likely to play an important role. Although also reprogrammed away from canonical exhaustion and into a superior effector state, endogenous tumor-specific TILs did not fully acquire the exact effector state of transferred cells, which presumably further benefitted from *in vitro* conditioning that was unavailable to natural T cells.

Tumor progression was not associated with evolution toward T-cell exhaustion or other kind of CD8 T-cell hyporesponsiveness states, but rather with extinction of the optimal effector states adopted by TILs. Confirming its dependence on the cooperative action of the two cytokines, loss of the optimal state coincided with attrition of IL-33 secreting cells. We attribute the loss of IL-33 producing cells to the decrease of the overall IL-2^V^ expression by the partner cells carrying the IL-2^V^ module, with likely decrease of its paracrine effects. Due to this imbalance, most residual engineered CD8^+^ TILs were arrested in less differentiated precursor-like states. Similarly, most endogenous cells converted to effector-memory, while residual Pex and Tex cells reacquired their initial TOX^+^ canonical exhausted phenotype, confirming that in the absence of both cytokines, these were the only available fates for TILs.

In summary, we provide proof-of-principle evidence that orthogonal combinatorial engineered CD8^+^ T cells for ACT, specifically secreting a variant of IL-2 that does not engage CD25 and the alarmin IL-33, can control advanced solid tumors, in the absence of preconditioning, cytokine therapy or other support (e.g. vaccination). While there are limited examples of ACT-mediated tumor control in lymphoreplete mice bearing hematological tumors using CAR-T cells (Avanzi et al. 2018; Kuhn et al. 2019), this represents to our knowledge the first such example of successful ACT in advanced, poorly immunogenic solid tumors in the absence of any supportive treatment. We employed a highly stringent solid tumor setting to demonstrate that orthogonal T-cell engineering can produce synthetic T cell states compatible with tumor eradication. The identification of the mechanism of tumor escape offers additional engineering opportunities, while the observation that CD4^+^ cell depletion enhanced the therapeutic effect points towards opportunities for additional combinatorial interventions targeting Tregs. Overall, our work demonstrates the potential for clinical translation of innovative combinatorial engineered T cells for reprogramming the TME in the immune competent host and inducing highly functional and novel synthetic CD8^+^ states endowed with the ability to control advanced, poorly immunogenic solid tumors.

## Acknowledgments

We thank Prof. Daniel Speiser and Werner Held for useful discussions and Dr. Kalet Leon (CIM, Havana, Cuba) for providing recombinant IL-2^V^ for in vitro T-cell expansion. We also thank Julien Marquis, Corinne Peter and Karolina Bojkowska from Lausanne Genomic Technologies Facility at UNIL for the support with the scRNAseq experiments.

## Author contributions

G.C., J.C.O. and M.I. conceived the project and designed the experiments; molecular cloning was performed by J.C.O. and B.S.; J.C.O., T.M., E.S., Y.O.M, W.C, C.J.L, A.S and C.R. performed experiments; E.L. optimized the retroviral transduction protocol. Analysis of scRNA-seq datasets was performed by J.C.O, S.J.C. and M.A.; histological analysis was designed and performed by S.A.L and L.S.; J.C.O., S.J.C. and G.C. interpreted data. The manuscript was written by G.C., J.C.O. P.R and M.I.; S.J.C., M.A and S.A.L. also contributed with the edition of the manuscript. The study was supervised by G.C. and M.I.

## Data and materials availability

All data is available from the authors upon reasonable request.

---

**Figure S1.**
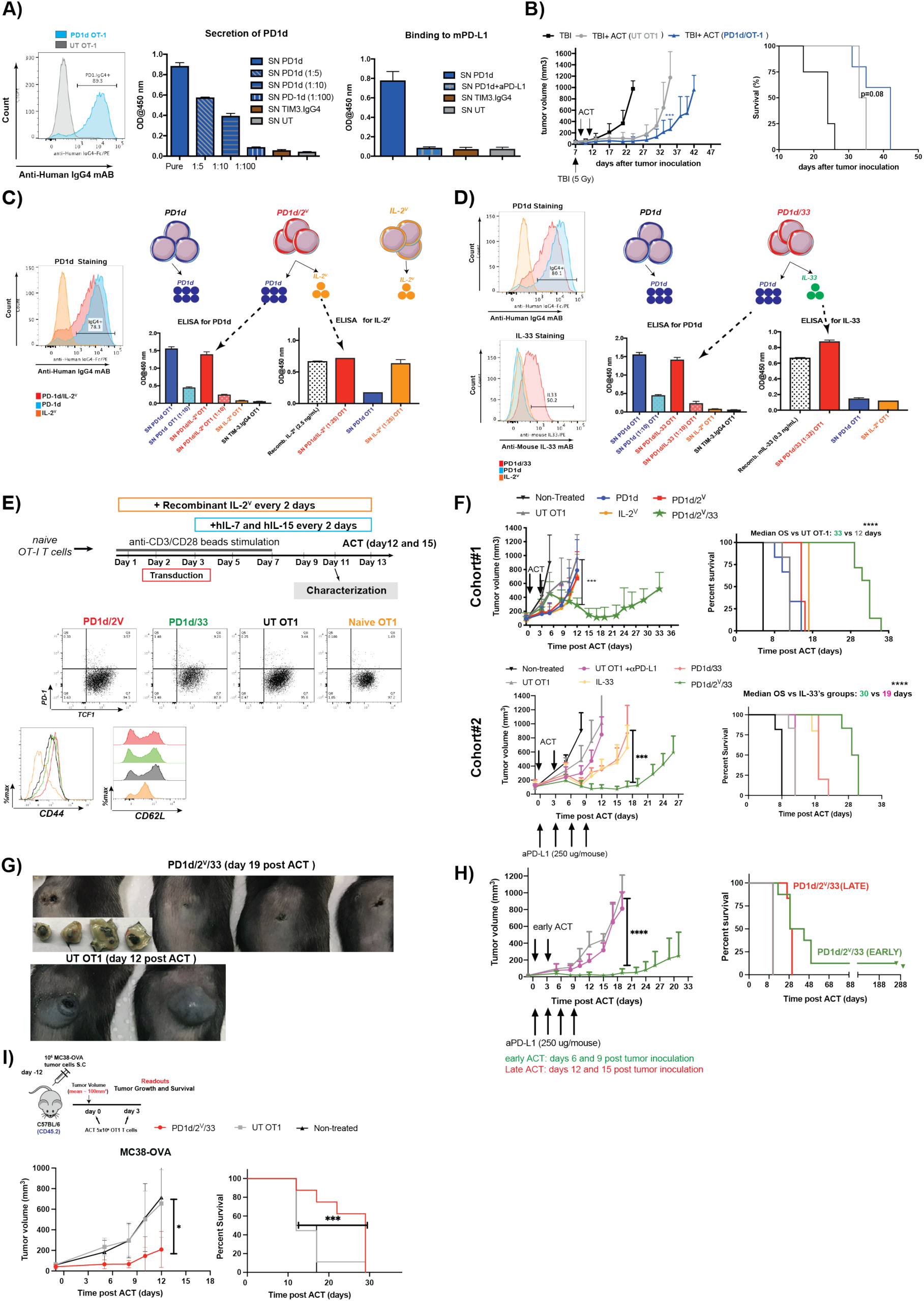
Relative to Figure1. **(A-D) Anti-tumor activity of OT-1 T cells gene-engineered for secreting the PD-1_IgG4 decoy in small B16-OVA tumors (∼ 30 mm^3^)**(**A**) Left: Intracellular expression of PD1d by FACS. Middle: Secretion into the SN determined by ELISA. Right: Binding to plate-coated mPD-L1. SN from OT-1 T cells gene-engineered to secrete TIM-3.IgG4 decoy was used as negative control. (**B**) Effect on tumor growth (left) and on overall survival (right) of ACT using OT1 cells gene-engineered to secrete the PD-1.IgG4 decoy. A representative experiment out of two independent experiments using 6 mice per group is shown. Survival Data were determined using a log-rank Mantel-Cox test. Tumor growth comparison at day 35 post tumor inoculation was done using a Mann-Whitney Test. *** p<0.001. (**C-D**) **OT-1 CD8 T cells can be gene-engineered to secrete PD1_IgG4 decoy in combination with either IL-2^V^ or IL-33** (**C**) Left: intracellular expression of PD1d in PD1d/2^V^ gene-engineered OT1 cells (there are no commercially available antibodies for detecting the IL-2^V^ using Flow Cytometry). Right: secretion of PD1d and IL-2^V^ into the supernatant by ELISA. (**D**) Left: Intracellular expression of PD1d and IL-33 in PD1d/33 gene-engineered OT1 cells. Right: secretion of PD1d and IL-33 into the supernatant by ELISA. OT1 cells gene-engineered to secrete TIM-3.IgG4 decoy were used as negative control to claim specificity in the PD-1_IgG4 ELISA. **(E) Phenotypic characterization of gene-engineered OT1 cells before ACT.** Top: schematic showing the transduction and expansion protocol of gene-engineered OT1 cells. Bottom: expression of CD44, TOX, CD62L, Ly6C and GzmC in expanded OT1 cells on day 10 post viral transduction. **(F) Orthogonal T cell engineering improves tumor control and survival in the absence of preconditioning and cytokine support**. Top: A representative cohort of mice treated with PD1d, IL-2^V^, PD1d/2^V^ and PD1d/2^V^/33. Bottom: a representative cohort of mice treated with UT OT1 plus αPD-L1, IL-33, PD1d/33 and PD1d/2^V^/33. Survival Data were determined using a log-rank Mantel-Cox test. Tumor growth comparison was done using a Mann-Whitney Test *** p<0.001, ****p<0.000. **(G)** Representative pictures of tumor control induced by PD1d/2^V^/33+OT1 or tumor progression of mice treated with untransduced OT1 (UT). (**H**) Tumor growth and survival curves of B16.OVA-bearing mice treated with either UT OT1 plus αPD-L1 or PD1d/2^V^/33+OT1 cells on days 6 and 9 after tumor cell inoculation. (**I**) Tumor growth and survival curves of MC38.OVA-bearing mice treated with either UT OT1 or PD1d/2^V^/33^+^OT1 cells on days 12 and 15 after tumor cell inoculation. Survival Data were determined using a log-rank Mantel-Cox test. Tumor growth comparison was done using a Mann-Whitney Test *** p<0.001, ****p<0.000

**Figure S2.**
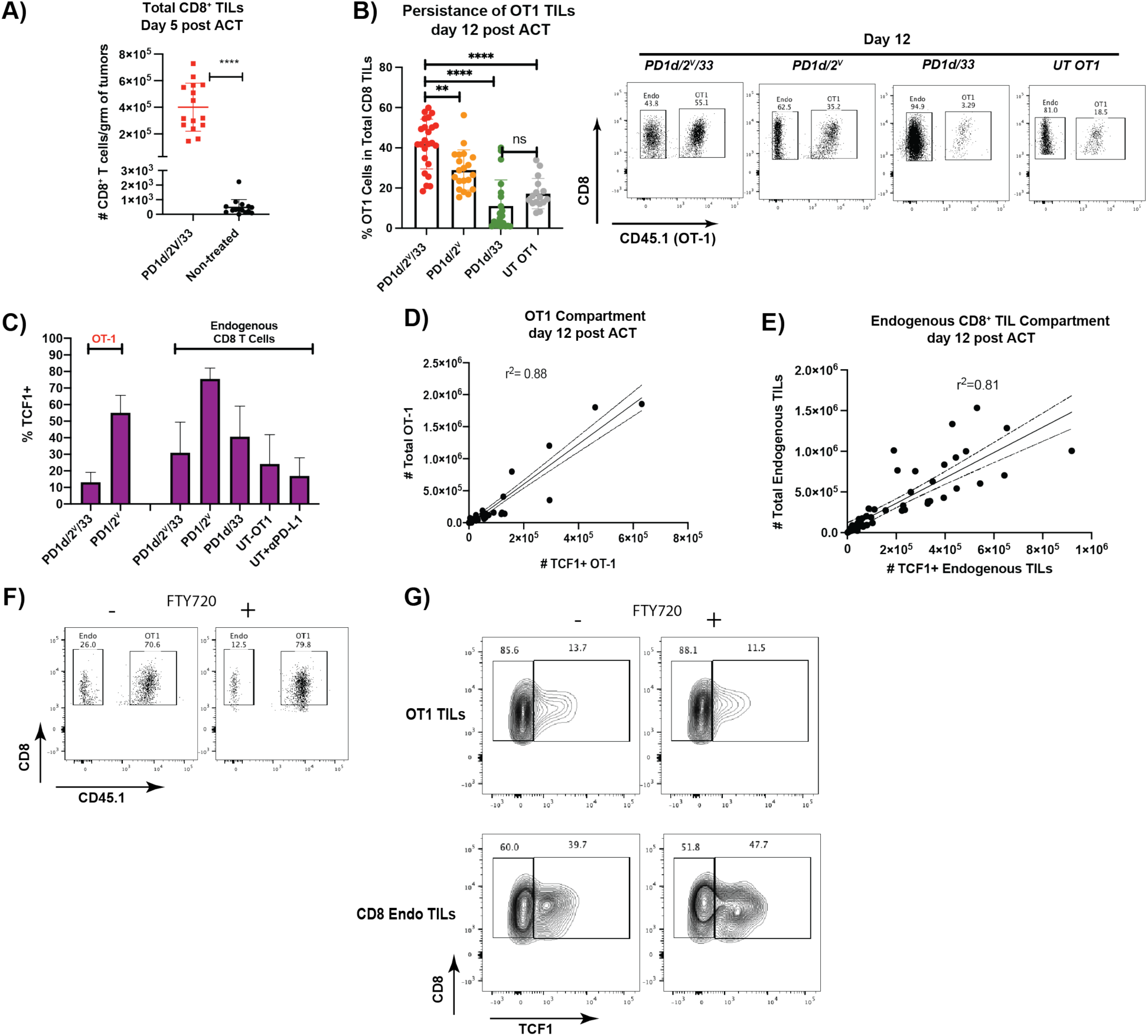
Relative to Figure1. Mice with B16.OVA tumors were treated with either engineered or untransduced OT1 cells on days 12 and 15 after tumor cell inoculation; then tumors were harvested on days 5 and 12 post ACT and cell quantification was performed by flow cytometry using precision counts beads. Data are from three independent experiments (n >= 5 animals/group). **(A)** Total numbers of CD8^+^ TILs at day 5 in mice treated with orthogonal ACT versus non-treated tumor bearing mice (17 days post tumor incoulation) evaluated in parallel. **(B)** Left: Intratumoral persistence of OT1 TILs at day 12 post ACT in each studied group. Right: representative dot plots for each treatment condition. **(C)** Analysis of the TCF1^+^ CD8^+^ TILs on day 12 post ACT. **(D)** Pearson correlation between the numbers of intratumoral TCF1^+^ OT1 T cells and the total number of intratumoral OT1 T cells on day 12 post orthogonal ACT **. (E)** Pearson correlation between the numbers of intratumoral TCF1^+^ endogenous CD8^+^ T cells and the total number of intratumoral endogenous CD8^+^ TILs at day 12 orthogonal ACT. **(F)** Representative dot plots showing the composition of the intratumoral CD8^+^ TIL compartment (Endogenous vs OT1 TILs) at day 12 post orthogonal ACT in the presence or absence of FTY720. **(G)** Representative contour plots showing the composition in TCF1^+^ and TCF1^neg^ TILs in both endogenous and OT1 TILs harvested at day 12 post orthogonal ACT in the presence or absence of FTY720.

**Figure S3.**
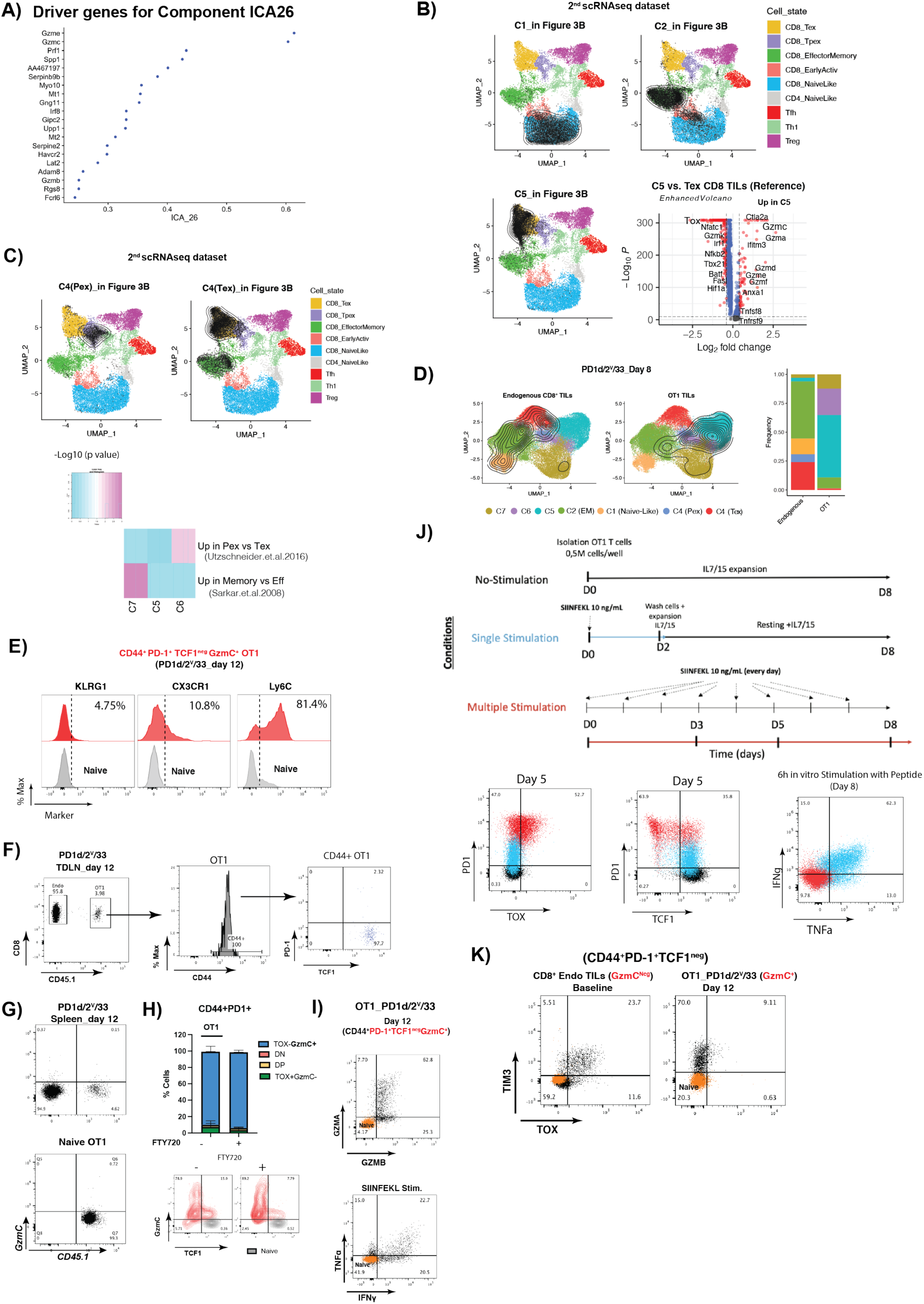
Relative to Figures 2 and 3. Complementary transcriptomic analysis of the different treatment conditions. (**A**) Driver Genes for the most significant independent component (ICA26) that separates cluster C5 from the TIL reference map. **(B)** Annotation of clusters C1-Naïve, C2-EffectorMemory and C5 from the 2^nd^ scRNAseq dataset based on their projection onto the reference mouse TIL map using ProjecTILs, where contour plots depict high cell density areas for each treatment. A volcano plot showing the differentially expressed genes between C5 cells and Tex cells from the reference TIL map is also shown. **(C)** Top: Annotation of clusters C4-Pex and C4-Tex from the 2^nd^ scRNAseq dataset based on their projection of on the reference mouse TIL map using ProjecTILs. Bottom: Gene-signature enrichment analysis against two reference CD8 T cell subtypes signatures observed in chronic and acute viral infections. (Sarkar et al. 2008), GEO accession GSE10239 and (Utzschneider et al. 2016), (GSE83978) respectively. Up refers to the sets of up-regulated genes associated with each comparison (e.g. Pex vs Tex UP: genes up-regulated in Pex). Color scale indicates statistical significance of signature overlap (FDR corrected *p*-values, Fisher’s exact test). **(D)** UMAP plot showing the cluster distribution of either OT1 or endogenous CD8 TILs recovered at day 8 post orthogonal ACT. Contour plots depict the clusters covered by each cell compartment. Bar plots are showing the cluster composition of each compartment. **(E)** Analysis of KLRG1, CX3CR1 and LY6C expression in CD44^+^ PD-1^+^ TCF1^neg^ GZMC^+^ OT1 TILs at day 12 post orthogonal ACT. **(F)** Analysis of CD44, PD-1 and expression in OT1 cells harvested at day 12 post ACT from the TDLN of mice treated with orthogonal ACT. **(G)** Analysis of GZMC expression in OT1 cells harvested at day 12 post ACT from the spleen of mice treated with orthogonal ACT. **(H)** Composition of the intratumoral CD8 TIL compartment (Endogenous vs OT1 TILs) at day 12 post orthogonal ACT in the presence or absence of FTY720 (**I)** Representative dot plots showing the expression of effector molecules in CD44^+^PD-1^+^TCF1^neg^GZMC^+^ OT1 TILs harvested at day 12 post orthogonal ACT. (**J)** Top: Schematics showing the experimental protocol to *in vitro* generate bona-fide exhausted OT1 T cells (Zhao et al. 2020). Bottom: Analysis of the expression of PD-1, TOX, TCF1, IFN and TNFα in multiple stimulated OT1 cells (red). OT1 cells that received one single stimulation (blue) or none (black), were used as controls. (**K**) Representative dot plots showing the expression of TIM3 and TOX in PD-1^+^ TCF1^neg^ Tex cells harvested at baseline (left) or in CD44^+^PD-1^+^TCF1^neg^GZMC^+^ OT1 TILs harvested at day 12 post orthogonal ACT (right)

**Figure S4.**
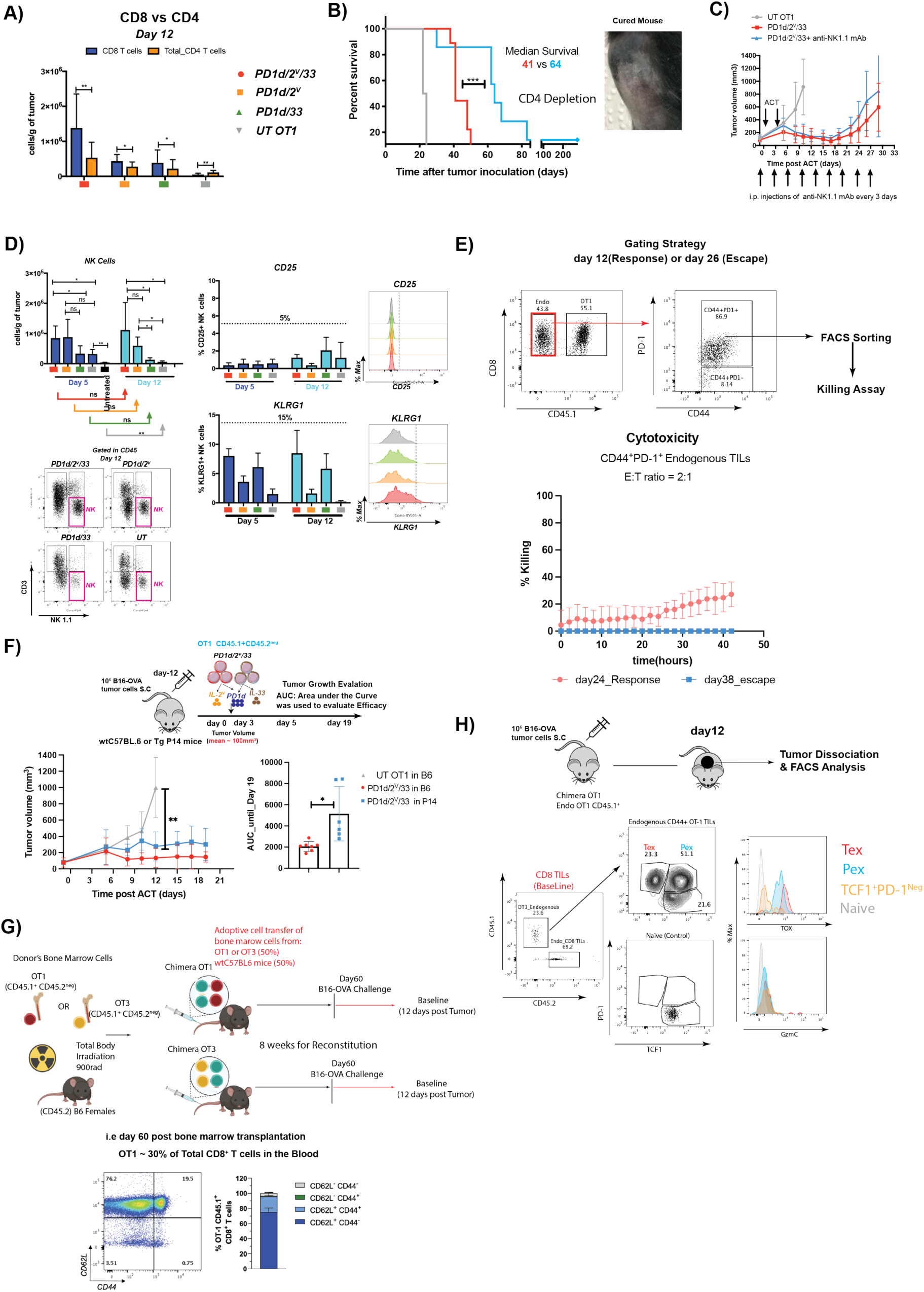
Relative to Figure 5. CD4 depletion significantly improves tumor control and survival of PD1d/2^V^/33-treated mice. **(A)** Tumor weight normalized numbers of CD4^+^ and CD8^+^ TILs on day 12 post ACT. **(B)** Survival over time curve of PD1d/2^V^/33+ OT1-treated mice in the absence or presence of a depleting antibody specific for CD4 administered i.p. 250 *μ*g/mouse beginning 1 d before initiation of PD1d/2^V^/33-ACT and maintained until day 43 post tumor inoculation. A representative picture of a cured mouse (more than a year free of tumor) is also shown. **(C-E**) **Intratumoral accumulation of NK Cells induced by orthogonal engineering is not linked to tumor control.** (**C**) Tumor growth over time curve of PD1d/2^V^/33+ OT1-treated mice in the absence or presence of a depleting antibody specific for NK1.1 administered i.p. 250 *μ*g/mouse beginning 1 d before initiation of PD1d/2^V^/33-ACT and maintained until day 27 post tumor inoculation. A Brown-Forsythe and Welch ANOVA test was used. Correction for multiple comparisons was done using a Tukey test. * p<0.05, ** p<0.01, *** p<0.001, ****p<0.0001 **(D)** Left: Tumor weight normalized numbers of NK cells on days 5 and 12 post ACT. Right: surface expression of NK-related activation markers: CD25 and KLRG1 on days 5 and 12 post ACT. (**E)** Top: gating strategy used to FACS sort endogenous CD8 TILs. Bottom: *In vitro* killing assay against B16-OVA tumors using Incucyte^TM^ of FACS-sorted CD44^+^PD-1^+^ endogenous CD8 TILs harvested at days 12 (Response, red) or day 26 (Escape, blue) post orthogonal ACT. **(F)** Top: Experimental design. Bottom left: PD1d/2^V^/33^+^ OT-1 cells were administrated as previously indicated, to B16.OVA-tumor bearing wt C57BL6 mice (red) or P14 transgenic mice (blue), (n = 6-7 mice group). Bottom right: area under the curve (AUC) for each individual mouse until day 19 post ACT (A representative experiment out of two independent experiments). A two-tailed Student’s t test with Welch’s correction was used for comparing the groups * p<0.05, ** p<0.01, *** p<0.001, ****p<0.0001. **(G)** Top: schematics summarizing the procedure to generate OT1 or OT3 mixed (50:50 with B6) bone marrow chimeras. Bottom: Phenotypic characterization of OT1 CD8 T cells harvested from the blood 2 months after bone marrow transplantation based on the expression of CD44 and CD62L. **(H)** Phenotypic characterization of endogenous OT1 TILs in B16.OVA tumor-bearing chimeric mice based on CD44, PD-1, TOX and GZMC expression.

**Figure S5.**
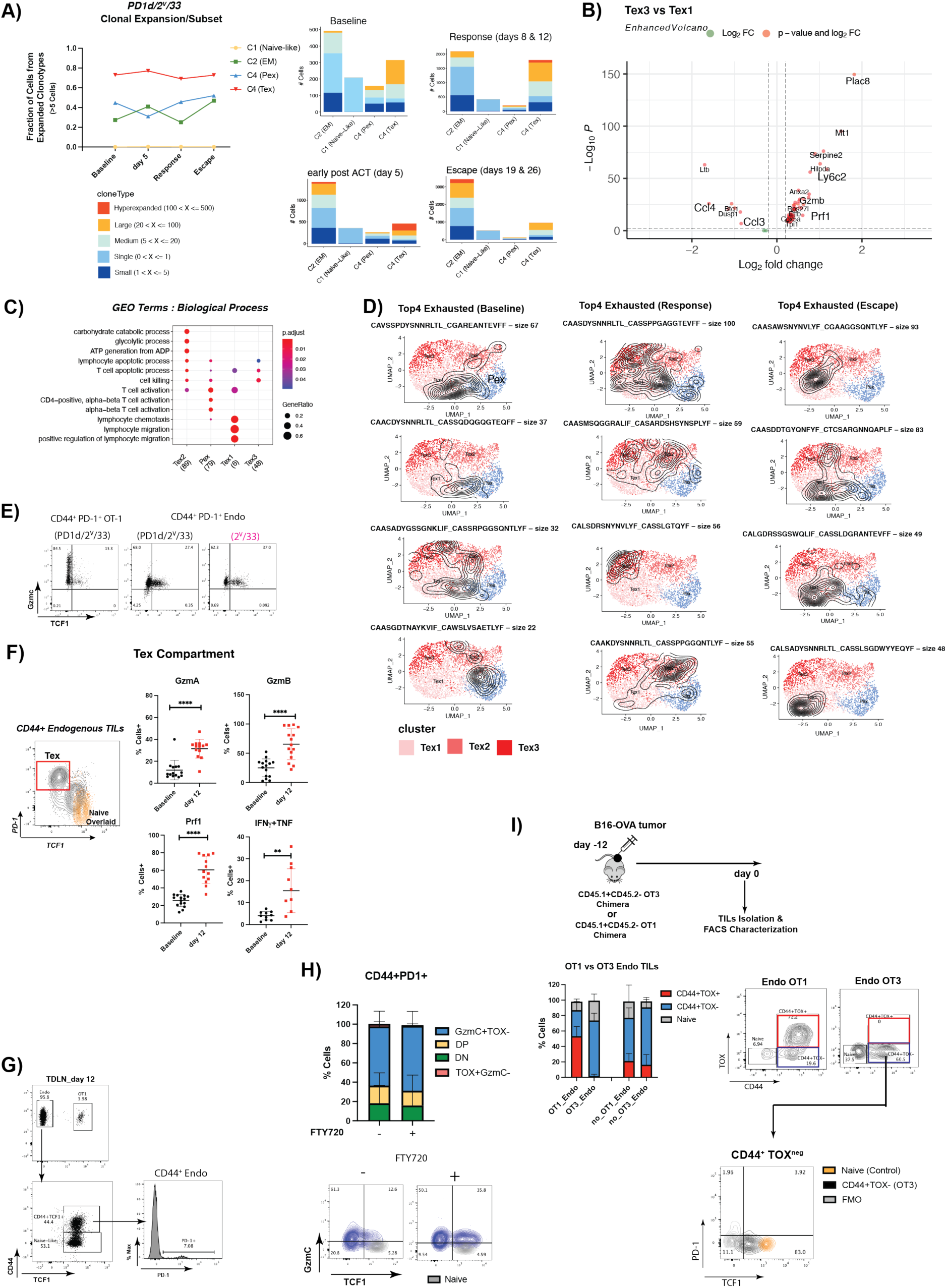
Relative to Figures 6. Orthogonal engineering reprograms endogenous Tex TILs. **(A)** Left: fraction of endogenous CD8 TILs by state that belong to expanded clonotypes (medium, large and hyperexpanded). Right: number of cells by state assigned into specific frequency ranges of clonal expansion in each experimental condition. **(B)** Volcano plot showing differentially expressed genes between Tex3 and Tex1 subtypes. **(C)** GEO terms enrichment analysis of each exhausted subtypes. **(D)** Distribution of the top 4 expanded exhausted clonotypes between Tex1, Tex2 and Tex3 subtypes in baseline tumors, or during the response or escape phases following orthogonal ACT. Contour plots depict the subtypes covered by each clonotype. **(E)** Representative dot plots showing expression analysis of GZMC and TCF1 in CD44^+^PD-1^+^ OT1 TILs harvested from responding mice upon orthogonal ACT (left part) or in CD44^+^PD-1^+^ endogenous Tex TILs from the same condition in the presence (middle) or absence (right) of PD1d. **(F)** Comparative functional analysis of PD-1^+^TCF1^neg^ Tex-like endogenous cells harvested at day 12 post orthogonal ACT versus Tex-like cells harvested from baseline tumors. **(G)** Analysis of CD44, PD-1 and TCF1 expression in endogenous CD8 T cells harvested at day 12 from the TDLN of mice treated with orthogonal ACT. **(H)** Analysis of TOX and GZMC expression in CD44^+^ PD-1^+^ Endogenous CD8^+^ TILs at day 12 post orthogonal ACT in the presence or absence of FTY720. **(I)** Top: experimental design and fraction of CD44^+^ TOX^+^ canonical exhausted TILs in endogenous OT1 versus OT3 from tumor-bearing chimeric mice. Bottom: Representative dot plots showing expression analysis of PD-1 and TCF1 in CD44^+^ TOX^neg^ endogenous OT3 TILs

**Figure S6.**
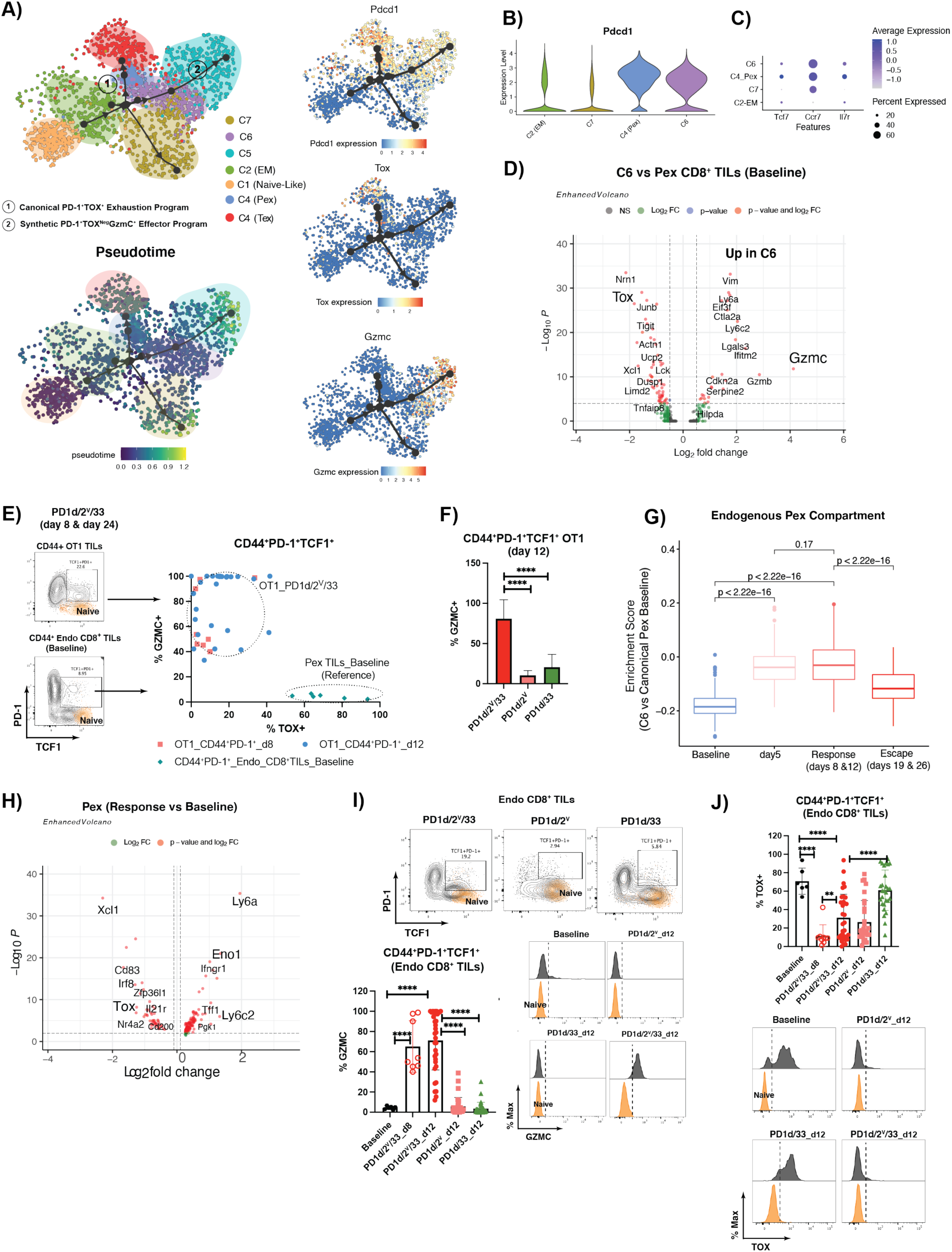
Orthogonally engineering ACT reprograms PD-1^+^ TCF1^+^ precursor-like TILs within the tumor microenvironment. **(A)** Trajectory and pseudotime inference analysis of OT1 TILs post harvested from all studied time points post orthogonal ACT using the Paga-Tree algorithm implemented in in *dynverse* (PMID: 30936559). The inferred trajectory as well as expression of relevant markers was visualized using diffusion maps. **(B)** Violin plots showing expression of *Pdcd1* gene in clusters C2-EM, C7, C4-Pex and C6. **(C)** Dot plot showing the expression of important stemness-related markers in clusters C2-EM, C7, C4-Pex and C6. **(D)** Comparison of C6 OT1 cells from days 8 and 12 (response) with canonical Pex-like cells recovered from baseline tumors. The volcano plot shows differentially expressed genes between both conditions. **(E)** Analysis of TOX and GZMC expression in PD-1**^+^**TCF1**^+^** OT1 TILs from orthogonal ACT relative to canonical Pex cells recovered from baseline tumors (n=6 mice). OT1 TILs from mice treated with orthogonal ACT were recovered at day 8 (two independent experiments n=4 mice/group) and at day 12 (5 independent experiments n=5-6 mice/group) animals/group) **(F)** Analysis of GZMC expression in CD44^+^ PD-1**^+^**TCF1^+^ OT1 TILs recovered at day 12 post ACT from mice treated with PD1d/2^V^, PD1d/33 or PD1d/2^V^/33, (data from 5 independent experiments n=5-6 mice/group). A representative dot plot is shown in the bottom part. One-way ANOVA test in combination with a Dunnet Test to correct for multiple comparisons was used * p<0.05, ** p<0.01, *** p<0.001, ****p<0.0001. **(G)** Enrichment of C6 gene signature (genes upregulated in C6 OT1 cells relative to baseline Pex) in endogenous Pex TILs harvested on days 8 and 12 from mice treated with PD1d/2^V^/33^+^OT1. The enrichment score of C6 gene signature was calculated with AUCell. A non-parametric Kruskal Wallis test was used for multiple comparisons * p<0.05, ** p<0.01, *** p<0.001, ****p<0.0001. **(H)** A volcano plot showing differentially expressed genes between Pex cells recovered from PD1d/2^V^/33-treated mice during the response phase (days 8 and 12) and Pex cells from baseline tumors. **(I-J)** Analysis of GZMC **(I)** and TOX **(J)** expression in CD44^+^ PD-1**^+^**TCF1^+^ endogenous TILs recovered at day 12 post ACT from mice treated with PD1d/2^V^, PD1d/33 and at days 8 and 12 post ACT from mice treated with PD1d/2^V^/33. Data collected from 5 independent experiments n=5-6 mice/group, except for samples collected at day 20 (2 independent experiments n=4 mice/group). One-way ANOVA test in combination with a Dunnet Test to correct for multiple comparisons was used * p<0.05, ** p<0.01, *** p<0.001, ****p<0.0001

**Figure S7.**
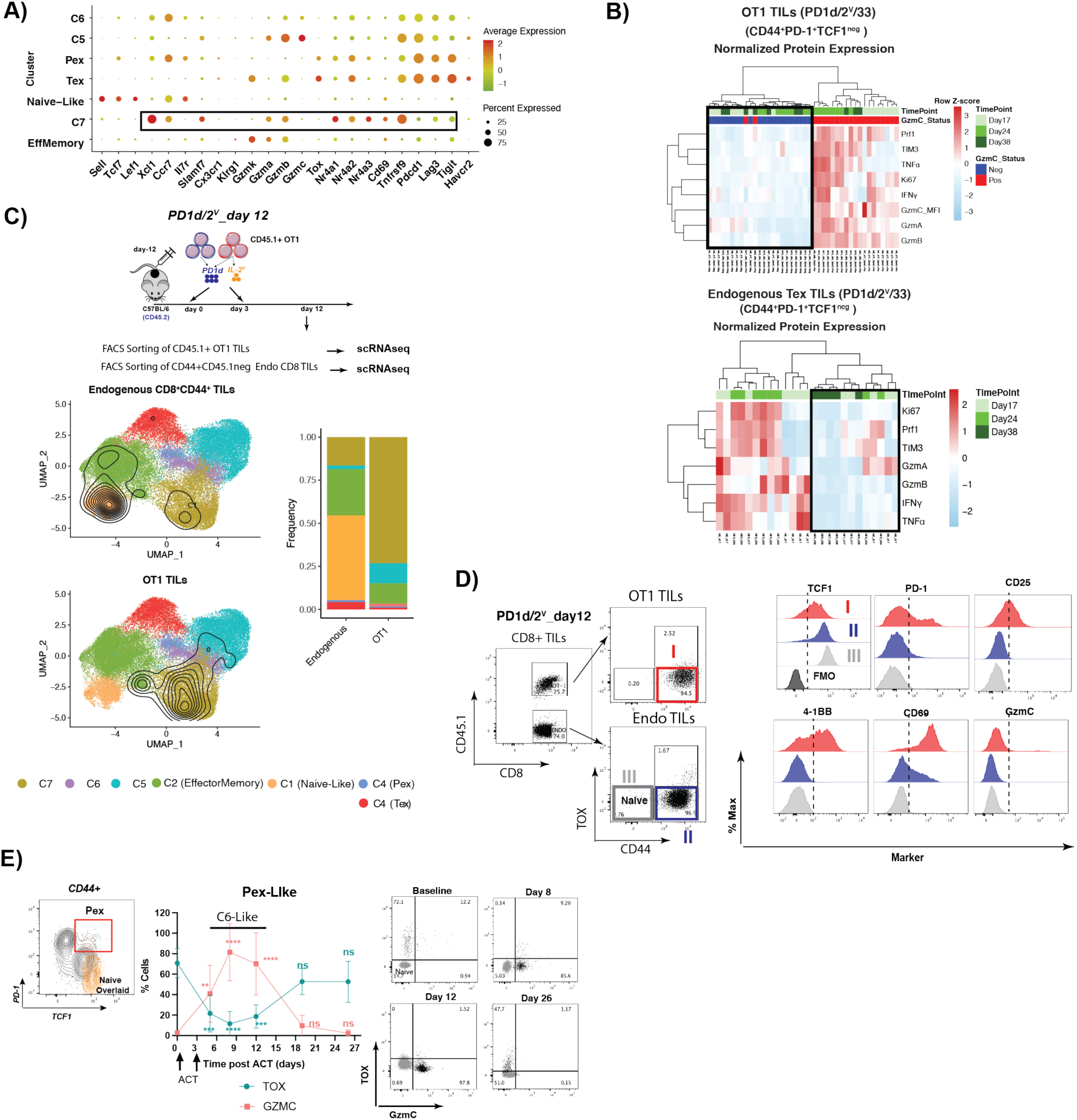
Relative to Figure 7. **(A)** Dot plot showing C7-specific markers **(B)** top: heatmap showing the normalized (row scaled) protein expression of TIM3, TNFα, GZMB, PRF1, IFN, GZMA, GZMB and KI67 in GZMC^+^ versus GZMC^neg^ CD44^+^ PD-1^+^ TCF1^neg^ OT1 TILs recovered at days 5, 12 and 26 post orthogonal ACT. Each column represents an individual mouse (n=19 mice in total). Bottom: like above but in endogenous Tex like cells. **(C)** UMAP plot showing the cluster distribution of either OT1 TILs or Endogenous CD8 TILs recovered at day 12 post ACT from mice treated with PD1d/2^V^ OT1. Contour plots depict the clusters covered by each cell compartment. Bar plots are showing the cluster composition of each compartment. **(D)** Phenotypic validation at the protein level of some C7-specific markers in either OT1 TILs or Endogenous CD8 TILs recovered at day 12 post ACT from mice treated with PD1d/2^V^ OT1. **(E)** Expression TOX and GZMC in endogenous Pex cells harvested at different time points following orthogonal ACT and defined as depicted in the dot plot based on the expression of TCF1 and PD-1. A two-tailed Student’s t test with Welch’s correction was used for comparing expression of each marker, in each time point relative to baseline Tex TILs * p<0.05, ** p<0.01, *** p<0.001, ****p<0.0001. Naïve CD8 T cells isolated from the spleen of non-tumor bearing mice were (gray) used as negative control for the expression of the studied markers.

**Table S1:**
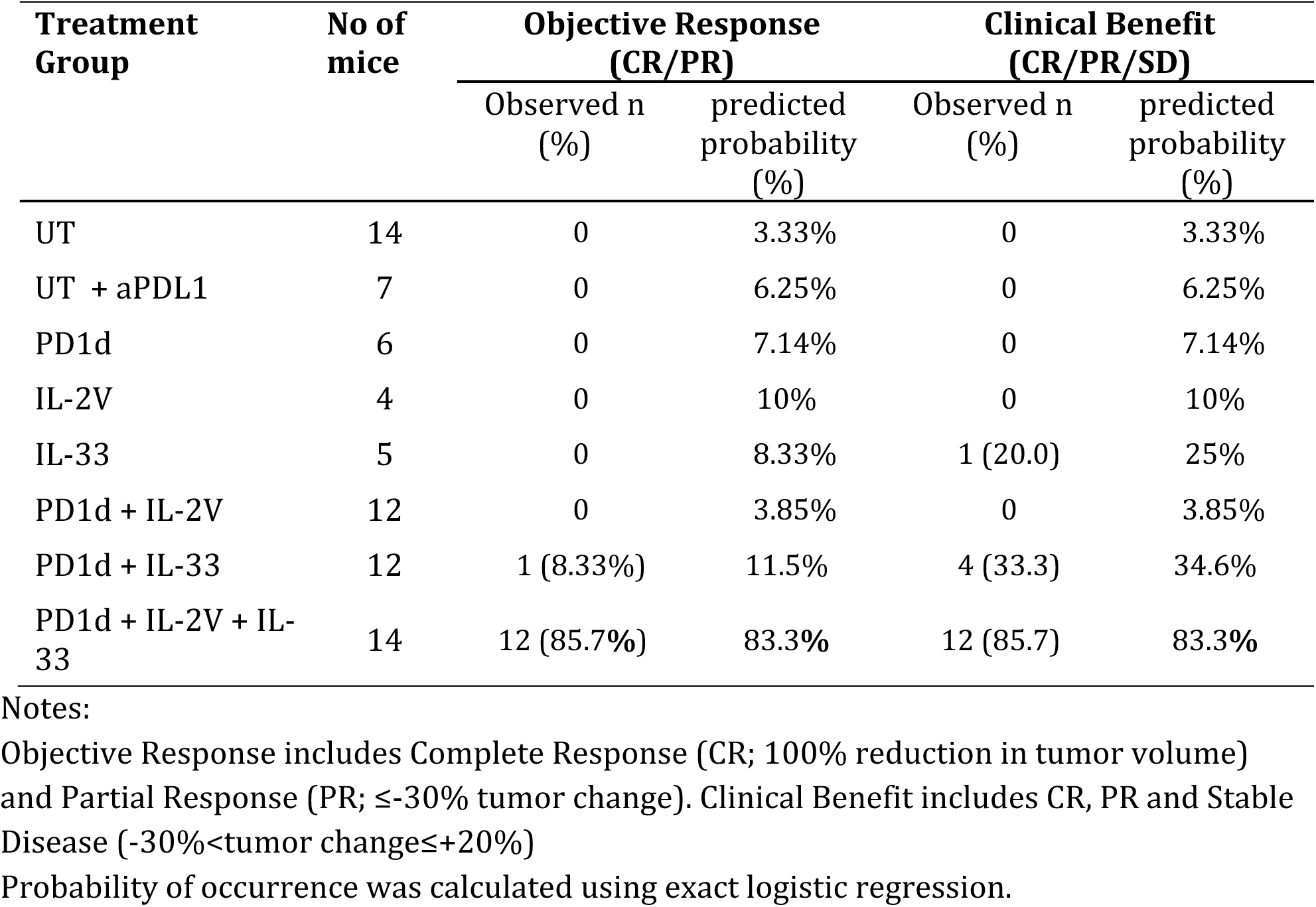
Observed response and predicted probability of objective response and clinical benefit for each treatment group.

| Treatment Group | No of mice | Objective Response (CR/PR) |  | Clinical Benefit (CR/PR/SD) |  |
| --- | --- | --- | --- | --- | --- |
|  |  | Observed n (%) | predicted probability (%) | Observed n (%) | predicted probability (%) |
| UT | 14 | 0 | 3.33% | 0 | 3.33% |
| UT + aPDL1 | 7 | 0 | 6.25% | 0 | 6.25% |
| PD1d | 6 | 0 | 7.14% | 0 | 7.14% |
| IL-2V | 4 | 0 | 10% | 0 | 10% |
| IL-33 | 5 | 0 | 8.33% | 1 (20.0) | 25% |
| PD1d + IL-2V | 12 | 0 | 3.85% | 0 | 3.85% |
| PD1d + IL-33 | 12 | 1 (8.33%) | 11.5% | 4 (33.3) | 34.6% |
| PD1d + IL-2V + IL-33 | 14 | 12 (85.7%) | 83.3% | 12 (85.7) | 83.3% |

Notes:
Objective Response includes Complete Response (CR; 100% reduction in tumor volume) and Partial Response (PR; $\leq -30\%$ tumor change). Clinical Benefit includes CR, PR and Stable Disease ( $-30\% < \text{tumor change} \leq +20\%$ )
Probability of occurrence was calculated using exact logistic regression.

**Table S2:** Predicted tumor size change from baseline for each group using linear regression.

| **Treatment** | **Predicted**<br>**(average) (%)**<br>**change in**<br>**tumor size** | **95% CI of the**<br>**expected (%)**<br>**change in tumor**<br>**size** | **p-value** |
| --- | --- | --- | --- |
| UT | 287.1% | (190% , 384%) | <0.0001 |
| UT + aPDL1 | 274.1% | (136% , 412%) | 0.0002 |
| PD1d | 343.7% | (195% , 493%) | <0.0001 |
| IL-2V | 608.0% | (426% , 790%) | <0.0001 |
| IL-33 | 145.7% | (-17% , 309%) | 0.0377 |
| PD1d + IL-2V | 290.8% | (186% , 396%) | <0.0001 |
| PD1d + IL-33 | 83.17% | (-22% , 188%) | 0.0570 |
| PD1d + IL-2V + IL-33 | -56% | (-100%* , 41%) | - |

\*: -100% is the minimum plausible value for tumor size change.
Notes: p-values compare each treatment effect versus the triple combination (PD1d + IL-2V + IL-33)
Adjusted R<sup>2</sup> of the model: 43.8%.

## References

1. Alfei, Francesca, Kristiyan Kanev, Maike Hofmann, Ming Wu, Hazem E. Ghoneim, Patrick Roelli, Daniel T. Utzschneider, Madlaina von Hoesslin, Jolie G. Cullen, Yiping Fan, Vasyl Eisenberg, Dirk Wohlleber, Katja Steiger, Doron Merkler, Mauro Delorenzi, Percy A. Knolle, Cyrille J. Cohen, Robert Thimme, Benjamin Youngblood, and Dietmar Zehn. 2019. ‘TOX reinforces the phenotype and longevity of exhausted T cells in chronic viral infection’, Nature, 571: 265–69.

2. Anderson, K. G., I. M. Stromnes, and P. D. Greenberg. 2017. ‘Obstacles Posed by the Tumor Microenvironment to T cell Activity: A Case for Synergistic Therapies’, Cancer Cell, 31: 311–25.

3. Andreatta, Massimo, Jesus Corria-Osorio, Sören Müller, Rafael Cubas, George Coukos, and Santiago J. Carmona. 2021. ‘Interpretation of T cell states from single-cell transcriptomics data using reference atlases’, Nature Communications, 12: 2965.

4. Avanzi, M. P., O. Yeku, X. Li, D. P. Wijewarnasuriya, D. G. van Leeuwen, K. Cheung, H. Park, T. J. Purdon, A. F. Daniyan, M. H. Spitzer, and R. J. Brentjens. 2018. ‘Engineered Tumor-Targeted T Cells Mediate Enhanced Anti-Tumor Efficacy Both Directly and through Activation of the Endogenous Immune System’, Cell Rep, 23: 2130–41.

5. Beltra, Jean-Christophe, Sasikanth Manne, Mohamed S. Abdel-Hakeem, Makoto Kurachi, Josephine R. Giles, Zeyu Chen, Valentina Casella, Shin Foong Ngiow, Omar Khan, Yinghui Jane Huang, Patrick Yan, Kito Nzingha, Wei Xu, Ravi K. Amaravadi, Xiaowei Xu, Giorgos C. Karakousis, Tara C. Mitchell, Lynn M. Schuchter, Alexander C. Huang, and E. John Wherry. 2020. ‘Developmental Relationships of Four Exhausted CD8+ T Cell Subsets Reveals Underlying Transcriptional and Epigenetic Landscape Control Mechanisms’, Immunity, 52: 825–41.e8.

6. Burger, Megan L., Amanda M. Cruz, Grace E. Crossland, Giorgio Gaglia, Cecily C. Ritch, Sarah E. Blatt, Arjun Bhutkar, David Canner, Tamina Kienka, Sara Z. Tavana, Alexia L. Barandiaran, Andrea Garmilla, Jason M. Schenkel, Michelle Hillman, Izumi de los Rios Kobara, Amy Li, Alex M. Jaeger, William L. Hwang, Peter M. K. Westcott, Michael P. Manos, Marta M. Holovatska, F. Stephen Hodi, Aviv Regev, Sandro Santagata, and Tyler Jacks. 2021. ‘Antigen dominance hierarchies shape TCF1+ progenitor CD8 T cell phenotypes in tumors’, Cell, 184: 4996–5014.e26.

7. Cai, Sheng F., Todd A. Fehniger, Xuefang Cao, Joshua C. Mayer, Joel D. Brune, Anthony R. French, and Timothy J. Ley. 2009. ‘Differential Expression of Granzyme B and C in Murine Cytotoxic Lymphocytes’, The Journal of Immunology, 182: 6287–97.

8. Carmenate, T., A. Pacios, M. Enamorado, E. Moreno, K. Garcia-Martinez, D. Fuente, and K. Leon. 2013. ‘Human IL-2 mutein with higher antitumor efficacy than wild type IL-2’, J Immunol, 190: 6230–8.

9. Carmona, Santiago J., Imran Siddiqui, Mariia Bilous, Werner Held, and David Gfeller. 2020. ‘Deciphering the transcriptomic landscape of tumor-infiltrating CD8 lymphocytes in B16 melanoma tumors with single-cell RNA-Seq’, OncoImmunology, 9: 1737369.

10. Chan, Jack D., Junyun Lai, Clare Y. Slaney, Axel Kallies, Paul A. Beavis, and Phillip K. Darcy. 2021. ‘Cellular networks controlling T cell persistence in adoptive cell therapy’, Nature Reviews Immunology, 21: 769–84.

11. Chen, Joyce, Isaac F. López-Moyado, Hyungseok Seo, Chan-Wang J. Lio, Laura J. Hempleman, Takashi Sekiya, Akihiko Yoshimura, James P. Scott-Browne, and Anjana Rao. 2019. ‘NR4A transcription factors limit CAR T cell function in solid tumours’, Nature, 567: 530–34.

12. Chen, Zeyu, Zhicheng Ji, Shin Foong Ngiow, Sasikanth Manne, Zhangying Cai, Alexander C. Huang, John Johnson, Ryan P. Staupe, Bertram Bengsch, Caiyue Xu, Sixiang Yu, Makoto Kurachi, Ramin S. Herati, Laura A. Vella, Amy E. Baxter, Jennifer E. Wu, Omar Khan, Jean-Christophe Beltra, Josephine R. Giles, Erietta Stelekati, Laura M. McLane, Chi Wai Lau, Xiaolu Yang, Shelley L. Berger, Golnaz Vahedi, Hongkai Ji, and E. John Wherry. 2019. ‘TCF-1-Centered Transcriptional Network Drives an Effector versus Exhausted CD8 T Cell-Fate Decision’, Immunity.

13. Chu, Talyn, and Dietmar Zehn. 2020. ‘Charting the Roadmap of T Cell Exhaustion’, Immunity, 52: 724–26.

14. Crompton, J. G., M. Sukumar, and N. P. Restifo. 2014. ‘Uncoupling T-cell expansion from effector differentiation in cell-based immunotherapy’, Immunol Rev, 257: 264–76.

15. Danilo, Maxime, Vijaykumar Chennupati, Joana Gomes Silva, Stefanie Siegert, and Werner Held. 2018. ‘Suppression of Tcf1 by Inflammatory Cytokines Facilitates Effector CD8 T Cell Differentiation’, Cell Rep, 22: 2107–17.

16. Dominguez, Donye, Cong Ye, Zhe Geng, Siqi Chen, Jie Fan, Lei Qin, Alan Long, Long Wang, Zhuoli Zhang, Yi Zhang, Deyu Fang, Timothy M. Kuzel, and Bin Zhang. 2017. ‘Exogenous IL-33 Restores Dendritic Cell Activation and Maturation in Established Cancer’, The Journal of Immunology, 198: 1365–75.

17. Feucht, Judith, Jie Sun, Justin Eyquem, Yu-Jui Ho, Zeguo Zhao, Josef Leibold, Anton Dobrin, Annalisa Cabriolu, Mohamad Hamieh, and Michel Sadelain. 2019. ‘Calibration of CAR activation potential directs alternative T cell fates and therapeutic potency’, Nat Med, 25: 82–88.

18. Gao, Kun, Xiaoying Li, Li Zhang, Lin Bai, Wei Dong, Kai Gao, Guiying Shi, Xianzhu Xia, Lingying Wu, and Lianfeng Zhang. 2013. ‘Transgenic expression of IL-33 activates CD8+ T cells and NK cells and inhibits tumor growth and metastasis in mice’, Cancer Letters, 335: 463–71.

19. Gao, Xin, Xuefeng Wang, Qianting Yang, Xin Zhao, Wen Wen, Gang Li, Junfeng Lu, Wenxin Qin, Yuan Qi, Fang Xie, Jingting Jiang, Changping Wu, Xueguang Zhang, Xinchun Chen, Heth Turnquist, Yibei Zhu, and Binfeng Lu. 2015. ‘Tumoral Expression of IL-33 Inhibits Tumor Growth and Modifies the Tumor Microenvironment through CD8^+^ T and NK Cells’, The Journal of Immunology, 194: 438–45.

20. Gattinoni, Luca, Steven E. Finkelstein, Christopher A. Klebanoff, Paul A. Antony, Douglas C. Palmer, Paul J. Spiess, Leroy N. Hwang, Zhiya Yu, Claudia Wrzesinski, David M. Heimann, Charles D. Surh, Steven A. Rosenberg, and Nicholas P. Restifo 2005. ‘Removal of homeostatic cytokine sinks by lymphodepletion enhances the efficacy of adoptively transferred tumor-specific CD8+ T cells’, Journal of Experimental Medicine, 202: 907–12.

21. Getachew, Yonas, Heather Stout-Delgado, Bonnie C. Miller, and Dwain L. Thiele. 2008. ‘Granzyme C Supports Efficient CTL-Mediated Killing Late in Primary Alloimmune Responses’, The Journal of Immunology, 181: 7810–17.

22. Held, Werner, Imran Siddiqui, Karin Schaeuble, and Daniel E. Speiser. 2019. ‘Intratumoral CD8^+^ T cells with stem cell–like properties: Implications for cancer immunotherapy’, Sci Transl Med, 11: eaay6863.

23. Hudson, William H., Julia Gensheimer, Masao Hashimoto, Andreas Wieland, Rajesh M. Valanparambil, Peng Li, Jian-Xin Lin, Bogumila T. Konieczny, Se Jin Im, Gordon J. Freeman, Warren J. Leonard, Haydn T. Kissick, and Rafi Ahmed. 2019. ‘Proliferating Transitory T Cells with an Effector-like Transcriptional Signature Emerge from PD-1+ Stem-like CD8+ T Cells during Chronic Infection’, Immunity, 51: 1043–58.e4.

24. Johnson, Hillary, Luca Scorrano, Stanley J. Korsmeyer, and Timothy J. Ley. 2003. ‘Cell death induced by granzyme C’, Blood, 101: 3093–101.

25. June, C. H., S. R. Riddell, and T. N. Schumacher. 2015. ‘Adoptive cellular therapy: a race to the finish line’, Sci Transl Med, 7: 280ps7.

26. Kalia, Vandana, Surojit Sarkar, Shruti Subramaniam, W. Nicholas Haining, Kendall A. Smith, and Rafi Ahmed. 2010. ‘Prolonged Interleukin-2R&#x3b1; Expression on Virus-Specific CD8^+^ T Cells Favors Terminal-Effector Differentiation In Vivo’, Immunity, 32: 91–103.

27. Kallert, Sandra M., Stephanie Darbre, Weldy V. Bonilla, Mario Kreutzfeldt, Nicolas Page, Philipp Müller, Matthias Kreuzaler, Min Lu, Stéphanie Favre, Florian Kreppel, Max Löhning, Sanjiv A. Luther, Alfred Zippelius, Doron Merkler, and Daniel D. Pinschewer. 2017. ‘Replicating viral vector platform exploits alarmin signals for potent CD8+ T cell-mediated tumour immunotherapy’, Nature Communications, 8: 15327.

28. Kallies, Axel, Dietmar Zehn, and Daniel T. Utzschneider. 2020. ‘Precursor exhausted T cells: key to successful immunotherapy?’, Nature Reviews Immunology, 20: 128–36.

29. Khan, Omar, Josephine R. Giles, Sierra McDonald, Sasikanth Manne, Shin Foong Ngiow, Kunal P. Patel, Michael T. Werner, Alexander C. Huang, Katherine A. Alexander, Jennifer E. Wu, John Attanasio, Patrick Yan, Sangeeth M. George, Bertram Bengsch, Ryan P. Staupe, Greg Donahue, Wei Xu, Ravi K. Amaravadi, Xiaowei Xu, Giorgos C. Karakousis, Tara C. Mitchell, Lynn M. Schuchter, Jonathan Kaye, Shelley L. Berger, and E. John Wherry. 2019. ‘TOX transcriptionally and epigenetically programs CD8+ T cell exhaustion’, Nature, 571: 211–18.

30. Klebanoff, C. A., S. A. Rosenberg, and N. P. Restifo. 2016. ‘Prospects for gene-engineered T cell immunotherapy for solid cancers’, Nat Med, 22: 26–36.

31. Kuhn, N. F., T. J. Purdon, D. G. van Leeuwen, A. V. Lopez, K. J. Curran, A. F. Daniyan, and R. J. Brentjens. 2019. ‘CD40 Ligand-Modified Chimeric Antigen Receptor T Cells Enhance Antitumor Function by Eliciting an Endogenous Antitumor Response’, Cancer Cell, 35: 473–88 e6.

32. Lynn, Rachel C., Evan W. Weber, Elena Sotillo, David Gennert, Peng Xu, Zinaida Good, Hima Anbunathan, John Lattin, Robert Jones, Victor Tieu, Surya Nagaraja, Jeffrey Granja, Charles F. A. de Bourcy, Robbie Majzner, Ansuman T. Satpathy, Stephen R. Quake, Michelle Monje, Howard Y. Chang, and Crystal L. Mackall. 2019. ‘c-Jun overexpression in CAR T cells induces exhaustion resistance’, Nature, 576: 293–300.

33. Mann, Thomas H., and Susan M. Kaech. 2019. ‘Tick-TOX, it’s time for T cell exhaustion’, Nat Immunol, 20: 1092–94.

34. Martinez, Gustavo J., Renata M. Pereira, Tarmo Äijö, Edward Y. Kim, Francesco Marangoni, Matthew E. Pipkin, Susan Togher, Vigo Heissmeyer, Yi Chen Zhang, Shane Crotty, Edward D. Lamperti, K. Mark Ansel, Thorsten R. Mempel, Harri Lähdesmäki, Patrick G. Hogan, and Anjana Rao. 2015. ‘The Transcription Factor NFAT Promotes Exhaustion of Activated CD8+ T Cells’, Immunity, 42: 265–78.

35. McLane, Laura M., Mohamed S. Abdel-Hakeem, and E. John Wherry. 2019. ‘CD8 T Cell Exhaustion During Chronic Viral Infection and Cancer’, Annual Review of Immunology, 37: 457–95.

36. Miller, Brian C., Debattama R. Sen, Rose Al Abosy, Kevin Bi, Yamini V. Virkud, Martin W. LaFleur, Kathleen B. Yates, Ana Lako, Kristen Felt, Girish S. Naik, Michael Manos, Evisa Gjini, Juhi R. Kuchroo, Jeffrey J. Ishizuka, Jenna L. Collier, Gabriel K. Griffin, Seth Maleri, Dawn E. Comstock, Sarah A. Weiss, Flavian D. Brown, Arpit Panda, Margaret D. Zimmer, Robert T. Manguso, F. Stephen Hodi, Scott J. Rodig, Arlene H. Sharpe, and W. Nicholas Haining. 2019. ‘Subsets of exhausted CD8+ T cells differentially mediate tumor control and respond to checkpoint blockade’, Nat Immunol, 20: 326–36.

37. Mognol, Giuliana P., Roberto Spreafico, Victor Wong, James P. Scott-Browne, Susan Togher, Alexander Hoffmann, Patrick G. Hogan, Anjana Rao, and Sara Trifari. 2017. ‘Exhaustion-associated regulatory regions in CD8^+^ tumor-infiltrating T cells’, Proceedings of the National Academy of Sciences, 114: E2776–E85.

38. Muranski, Pawel, Andrea Boni, Claudia Wrzesinski, Deborah E. Citrin, Steven A. Rosenberg, Richard Childs, and Nicholas P. Restifo. 2006. ‘Increased intensity lymphodepletion and adoptive immunotherapy—how far can we go?’, Nature Clinical Practice Oncology, 3: 668–81.

39. Obar, Joshua J., Michael J. Molloy, Evan R. Jellison, Thomas A. Stoklasek, Weijun Zhang, Edward J. Usherwood, and Leo Lefrançois. 2010. ‘CD4^+^ T cell regulation of CD25 expression controls development of short-lived effector CD8^+^ T cells in primary and secondary responses’, Proceedings of the National Academy of Sciences, 107: 193–98.

40. Pipkin, Matthew E., Jilian A. Sacks, Fernando Cruz-Guilloty, Mathias G. Lichtenheld, Michael J. Bevan, and Anjana Rao. 2010. ‘Interleukin-2 and Inflammation Induce Distinct Transcriptional Programs that Promote the Differentiation of Effector Cytolytic T Cells’, Immunity, 32: 79–90.

41. Rojas, Gertrudis, Tania Carmenate, Julio Felipe Santo-Tomás, Pedro A. Valiente, Marlies Becker, Annia Pérez-Riverón, Yaima Tundidor, Yaquelín Ortiz, Jorge Fernandez de Cossio-Diaz, Luis Graça, Stefan Dübel, and Kalet León. 2019. ‘Directed evolution of super-secreted variants from phage-displayed human Interleukin-2’, Scientific Reports, 9: 800.

42. Rosenberg, S. A., and N. P. Restifo. 2015. ‘Adoptive cell transfer as personalized immunotherapy for human cancer’, Science, 348: 62–8.

43. Sarkar, Surojit, Vandana Kalia, W. Nicholas Haining, Bogumila T. Konieczny, Shruti Subramaniam, and Rafi Ahmed 2008. ‘Functional and genomic profiling of effector CD8 T cell subsets with distinct memory fates’, Journal of Experimental Medicine, 205: 625–40.

44. Scott, Andrew C., Friederike Dündar, Paul Zumbo, Smita S. Chandran, Christopher A. Klebanoff, Mojdeh Shakiba, Prerak Trivedi, Laura Menocal, Heather Appleby, Steven Camara, Dmitriy Zamarin, Tyler Walther, Alexandra Snyder, Matthew R. Femia, Elizabeth A. Comen, Hannah Y. Wen, Matthew D. Hellmann, Niroshana Anandasabapathy, Yong Liu, Nasser K. Altorki, Peter Lauer, Olivier Levy, Michael S. Glickman, Jonathan Kaye, Doron Betel, Mary Philip, and Andrea Schietinger. 2019. ‘TOX is a critical regulator of tumour-specific T cell differentiation’, Nature, 571: 270–74.

45. Seo, Hyungseok, Joyce Chen, Edahí González-Avalos, Daniela Samaniego-Castruita, Arundhoti Das, Yueqiang H. Wang, Isaac F. López-Moyado, Romain O. Georges, Wade Zhang, Atsushi Onodera, Cheng-Jang Wu, Li-Fan Lu, Patrick G. Hogan, Avinash Bhandoola, and Anjana Rao. 2019. ‘TOX and TOX2 transcription factors cooperate with NR4A transcription factors to impose CD8^+^ T cell exhaustion’, Proceedings of the National Academy of Sciences, 116: 12410–15.

46. Sheih, Alyssa, Valentin Voillet, Laïla-Aïcha Hanafi, Hannah A. DeBerg, Masanao Yajima, Reed Hawkins, Vivian Gersuk, Stanley R. Riddell, David G. Maloney, Martin E. Wohlfahrt, Dnyanada Pande, Mark R. Enstrom, Hans-Peter Kiem, Jennifer E. Adair, Raphaël Gottardo, Peter S. Linsley, and Cameron J. Turtle. 2020. ‘Clonal kinetics and single-cell transcriptional profiling of CAR-T cells in patients undergoing CD19 CAR-T immunotherapy’, Nature Communications, 11: 219.

47. Siddiqui, Imran, Karin Schaeuble, Vijaykumar Chennupati, Silvia A. Fuertes Marraco, Sandra Calderon-Copete, Daniela Pais Ferreira, Santiago J. Carmona, Leonardo Scarpellino, David Gfeller, Sylvain Pradervand, Sanjiv A. Luther, Daniel E. Speiser, and Werner Held. 2019. ‘Intratumoral Tcf1+PD-1+CD8+ T Cells with Stem-like Properties Promote Tumor Control in Response to Vaccination and Checkpoint Blockade Immunotherapy’, Immunity, 50: 195–211.e10.

48. Slade, Chris D., Katie L. Reagin, Hari G. Lakshmanan, Kimberly D. Klonowski, and Wendy T. Watford. 2020. ‘Placenta-specific 8 limits IFNγ production by CD4 T cells in vitro and promotes establishment of influenza-specific CD8 T cells in vivo’, PLOS ONE, 15: e0235706.

49. Utzschneider, Daniel T., Mélanie Charmoy, Vijaykumar Chennupati, Laurène Pousse, Daniela Pais Ferreira, Sandra Calderon-Copete, Maxime Danilo, Francesca Alfei, Maike Hofmann, Dominik Wieland, Sylvain Pradervand, Robert Thimme, Dietmar Zehn, and Werner Held. 2016. ‘T Cell Factor 1-Expressing Memory-like CD8+ T Cells Sustain the Immune Response to Chronic Viral Infections’, Immunity, 45: 415–27.

50. Villarreal, Daniel O., and David B. Weiner. 2014. ‘Interleukin 33: a switch-hitting cytokine’, Curr Opin Immunol, 28: 102–06.

51. Villarreal, Daniel O., Megan C. Wise, Jewell N. Walters, Emma L. Reuschel, Min Joung Choi, Nyamekye Obeng-Adjei, Jian Yan, Matthew P. Morrow, and David B. Weiner. 2014. ‘Alarmin IL-33 Acts as an Immunoadjuvant to Enhance Antigen-Specific Tumor Immunity’, Cancer Research, 74: 1789–800.

52. Vodnala, Suman Kumar, Robert Eil, Rigel J. Kishton, Madhusudhanan Sukumar, Tori N. Yamamoto, Ngoc-Han Ha, Ping-Hsien Lee, MinHwa Shin, Shashank J. Patel, Zhiya Yu, Douglas C. Palmer, Michael J. Kruhlak, Xiaojing Liu, Jason W. Locasale, Jing Huang, Rahul Roychoudhuri, Toren Finkel, Christopher A. Klebanoff, and Nicholas P. Restifo. 2019. ‘T cell stemness and dysfunction in tumors are triggered by a common mechanism’, Science, 363: eaau0135.

53. Wolf, Benita, Stefan Zimmermann, Caroline Arber, Melita Irving, Lionel Trueb, and George Coukos. 2019. ‘Safety and Tolerability of Adoptive Cell Therapy in Cancer’, Drug Safety, 42: 315–34.

54. Yao, Chen, Hong-Wei Sun, Neal E. Lacey, Yun Ji, E. Ashley Moseman, Han-Yu Shih, Elisabeth F. Heuston, Martha Kirby, Stacie Anderson, Jun Cheng, Omar Khan, Robin Handon, Julie Reilley, Jessica Fioravanti, Jinhui Hu, Selamawit Gossa, E. John Wherry, Luca Gattinoni, Dorian B. McGavern, John J. O’Shea, Pamela L. Schwartzberg, and Tuoqi Wu. 2019. ‘Single-cell RNA-seq reveals TOX as a key regulator of CD8+ T cell persistence in chronic infection’, Nat Immunol, 20: 890–901.

55. Yu, Jong W., Sabyasachi Bhattacharya, Niranjan Yanamandra, David Kilian, Hong Shi, Sapna Yadavilli, Yuliya Katlinskaya, Heather Kaczynski, Michael Conner, William Benson, Ashleigh Hahn, Laura Seestaller-Wehr, Meixia Bi, Nicholas J. Vitali, Lyuben Tsvetkov, Wendy Halsey, Ashley Hughes, Christopher Traini, Hui Zhou, Junping Jing, Tae Lee, David J. Figueroa, Sara Brett, Christopher B. Hopson, James F. Smothers, Axel Hoos, and Roopa Srinivasan. 2018. ‘Tumor-immune profiling of murine syngeneic tumor models as a framework to guide mechanistic studies and predict therapy response in distinct tumor microenvironments’, PLOS ONE, 13: e0206223.

56. Zander, Ryan, David Schauder, Gang Xin, Christine Nguyen, Xiaopeng Wu, Allan Zajac, and Weiguo Cui. 2019. ‘CD4+ T Cell Help Is Required for the Formation of a Cytolytic CD8+ T Cell Subset that Protects against Chronic Infection and Cancer’, Immunity, 51: 1028–42.e4.

57. Zhao, M., C. H. Kiernan, C. J. Stairiker, J. L. Hope, L. G. Leon, M. van Meurs, I. Brouwers-Haspels, R. Boers, J. Boers, J. Gribnau, IJcken W. F. J. van, E. M. Bindels, R. M. Hoogenboezem, S. J. Erkeland, Y. M. Mueller, and P. D. Katsikis. 2020. ‘Rapid in vitro generation of bona fide exhausted CD8+ T cells is accompanied by Tcf7 promotor methylation’, PLoS Pathog, 16: e1008555.

58. Zhao, Xudong, Qiang Shan, and Hai-Hui Xue. 2021. ‘TCF1 in T cell immunity: a broadened frontier’, Nature Reviews Immunology.

